# Integrated *de novo* Gene Prediction and Peptide Assembly of Metagenomic Sequencing Data

**DOI:** 10.1101/2021.09.20.461079

**Authors:** Sirisha Thippabhotla, Ben Liu, Shibu Yooseph, Youngik Yang, Jun Zhang, Cuncong Zhong

## Abstract

Metagenomics is the study of all genomic content presented in given microbial communities. Metagenomic functional analysis aims to quantify protein families and reconstruct metabolic pathways from the metagenome. It plays a central role in understanding the interaction between the microbial community and its host or environment. *De novo* functional analysis, which allows the discovery of novel protein families, remains challenging for high-complexity communities. There are currently three main approaches for recovering novel genes or proteins: *de novo* nucleotide assembly, gene calling, and peptide assembly. Unfortunately, their informational connection and dependency have been overlooked, and each has been formulated as an independent problem. In this work, we develop a sophisticated workflow called integrated Metagenomic Protein Predictor (iMPP), which leverages the informational dependencies for better *de novo* functional analysis. iMPP contains three novel modules: a hybrid assembly graph generation module, a graph-based gene calling module, and a peptide assembly-based refinement module. iMPP significantly improved the existing gene calling sensitivity on unassembled fragmented reads, achieving a 92% - 97% recall rate at a high precision level (>90%). iMPP further allowed for more sensitive and accurate peptide assembly, recovering more reference proteins and delivering more hypothetical protein sequences. The high performance of iMPP can provide a more comprehensive and unbiased view of the microbial communities under investigation. iMPP is freely available from https://github.com/Sirisha-t/iMPP.

## INTRODUCTION

Microbial communities are ubiquitously present in many environmental niches on earth, including soil (1), water (2), and air (3). Microbial communities are also a critical component of the human system, playing important roles in maintaining human health and wellbeing (4-6). Human microbiome dysbiosis can lead to various diseases, such as obesity (7-10), diabetes (11,12), and inflammatory bowel disease (13-15). On the other hand, human microbiome intervention has recently been explored as a meaningful non-invasive treatment. For example, *Salmonella, Escherichia*, and *Clostridium* are used as anticancer agents with highly promising effects in cancer therapeutics (16-18). Certain microbes also correlate with the response and toxicity from cancer treatments (19,20). Advances in next-generation sequencing (NGS) enable the study of the genomic content of a microbial community as a whole, known as metagenomics (21,22). Metagenomic sequencing data allows one to examine the taxonomic composition of the microbial community (23-25). More importantly, it further enables protein family profiling (26-28) and metabolic pathway reconstruction (29,30). This information is critical to unlocking the functional potential of the microbial community and elucidating its interactions with the environment.

Metagenomic functional analysis usually begins with homology search, such as aligning the sequencing reads against functionally annotated genomes (e.g., NCBI RefSeq) or protein databases (NCBI NR or UniProt (31)) using BLAST (32). However, due to the incompleteness of current databases, this approach may overlook functional elements encoded by previously-unseen microbial species and novel protein families, yielding a biased view of the community’s function. Alternatively, a reference-independent approach first assembles the sequencing reads into complete or near-complete genome sequences using *de novo* genome assemblers such as Meta-IDBA (33), MEGAHIT (34), MetaVelvet (35,36), and metaSPAdes (37). Then, it attempts to find open reading frames (ORFs) directly from the assembled genomes based on signals such as gene length, GC-content, and codon usage that are universal among all protein-coding genes. The so-called *de novo* gene calling step can be handled by software packages like Glimmer (38), GeneMark (39), and Prodigal (40). When long enough genomic sequences with stable and complete ORF signals are available, *de novo* gene calling is often reliable. However, the problem becomes more challenging on fragmented sequences (e.g., unassembled reads). More sophisticated computational models and algorithms are often required to solve the problem. Software packages that support fragmented gene calling include MetaGeneAnnotator (41), FragGeneScan (42), Orphelia (43), Glimmer-MG (44), MetaGeneMark (45), and MetaProdigal (46). Despite being less accurate than their genome-scale counterparts (42,43), fragmented gene callers can detect low-abundance protein-coding reads that are difficult to assemble. They output the detected protein-coding reads, whose corresponding peptide sequences can be further assembled into peptide contigs using *de novo* peptide assemblers such as SPA (47,48), PLASS (49), and MetaPA (50).

The three *de novo* functional analysis approaches discussed above, i.e., *de novo* nucleotide assembly, gene calling, and peptide assembly, strongly depend on each other (Figure 1). First, nucleotide assembly reconstructs longer genomic sequences with stronger and more stable ORF signals, which is expected to improve gene calling (42,43). For example, Graph2Pro (51) explicitly couples nucleotide assembly and gene calling by searching ORFs from paths in the nucleotide assembly graph. Second, gene calling can benefit downstream peptide assembly by providing refined short peptide sequences as input. The peptide assembler SPA (47) showed a higher performance when fed with peptide sequences predicted by FragGeneScan (42) compared to those predicted by MetaGeneAnnotator (41). We will further show (in this work) that peptide assemblers that accept all six-frame translations as input can also benefit from a refined input set. Conversely, peptide assembly explores the overlap information among the input short peptide sequences and can improve gene calling by rescuing false-negative predictions. Specifically, if a candidate ORF significantly overlaps with other peptides and is assembled into a long-enough contig, the candidate ORF is likely to be correct. The peptide overlap information is independent of the traditional ORF signals (e.g., codon frequency) and can further contribute to gene calling. Finally, peptide assembly reconstructs longer peptide contigs or even complete protein sequences that can serve as guides to nucleotide assembly. The so-called gene-centric assembly demonstrates better performance than its model-free counterparts (52-54).

**Figure 1:**
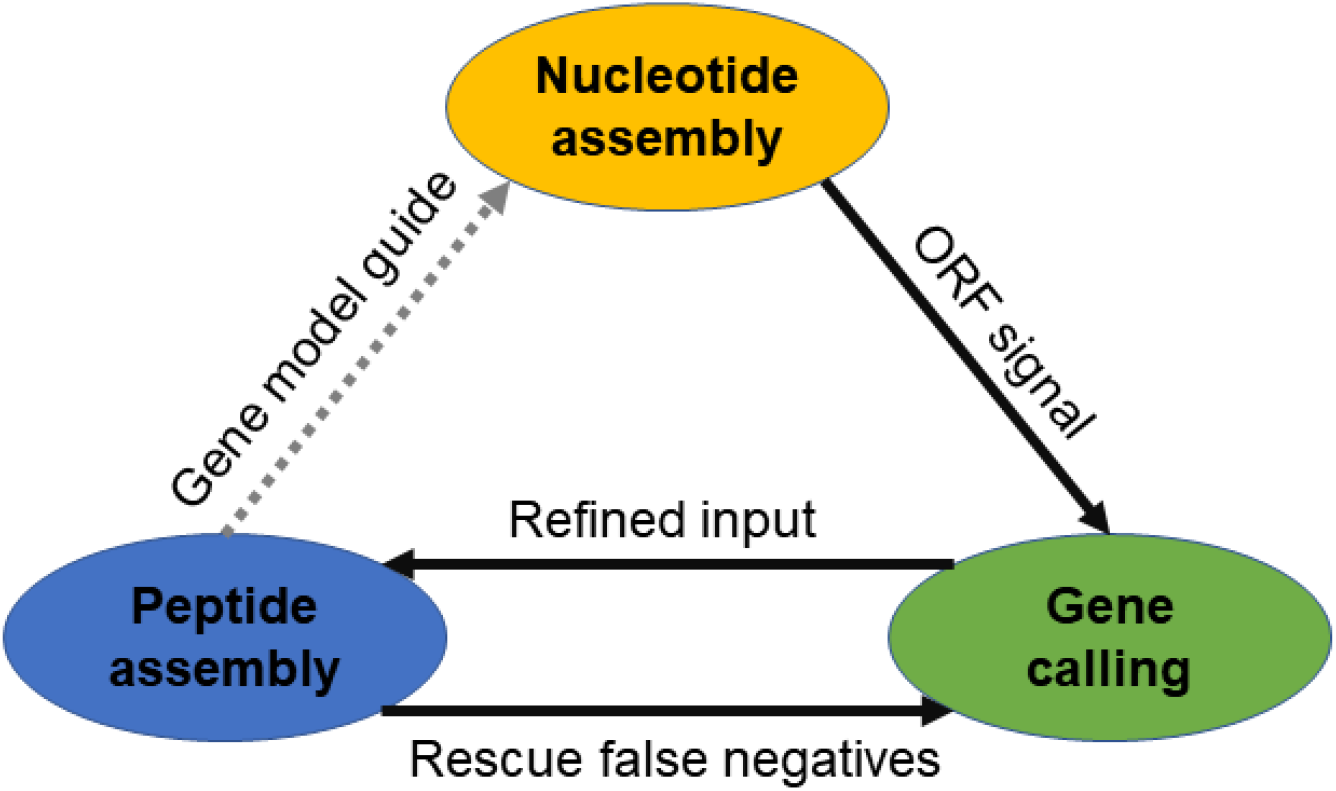
The informational dependency among de novo nucleotide assembly, gene calling, and peptide assembly. Solid arrows indicate dependencies that have been utilized by iMPP to improve metagenomic functional annotation.

Despite the strong informational connection and dependency of *de novo* nucleotide assembly, gene calling, and peptide assembly in metagenomic functional analysis, they have largely been considered and solved independently. Examples include many dedicated metagenome assemblers (33-37), dedicated metagenomic gene callers (41,42,44,45), and dedicated metagenomic peptide assemblers (47-50). While Graph2Pro (51) explicitly couples nucleotide assembly with gene calling, it expects metaproteomic data to validate its protein prediction and lacks a peptide assembly component. To the best of our knowledge, no functional annotation method exists that considers the informational dependency among the three approaches and integrates them into a single functional analysis framework. It remains unclear whether doing so is feasible and by how much it can improve metagenomic functional analysis.

We integrate nucleotide assembly, gene calling, and peptide assembly into a *de novo* metagenomic functional analysis workflow called integrated Metagenomic Protein Predictor (iMPP). Instead of being a simple sequential execution, iMPP is empowered with three novel modules to fully leverage the informational dependency. iMPP constructs a hybrid assembly graph by merging a de Bruijn graph and an overlap graph. The de Bruijn graph information increases graph connectedness, while the overlap graph information retains minor sequence variations. It further contains a novel gene calling module that operates on the merged hybrid graph. The gene calling module is computationally efficient by applying heuristics to eliminate unnecessary graph traversals. Finally, iMPP employs a protein reconstruction module with a two-pass peptide assembly, correcting the gene calling results in the first pass and reconstructing peptide contigs in the second pass. Due to computational efficiency concerns, the current implementation of iMPP does not contain a gene-centric nucleotide assembly module that guides nucleotide assembly with the assembled peptide contigs (Figure 1, the broken gray line).

We benchmarked the performance of iMPP in terms of both *de novo* gene calling and peptide sequence assembly on four real metagenomic datasets from different environments: human gut, soil, marine, and cow rumen. While the performance of the existing gene calling methods is already as high as 85-90%, iMPP further improved it by another ∼5%, reaching ∼92-93% of F-measure. For peptide assembly, we further compared iMPP with two other strategies: one as a sequential integration of gene calling and peptide assembly, and the other as peptide assembly alone. Our evaluations using both real and simulated metagenomic datasets showed that iMPP outperformed both strategies in most assembly statistics, including assembly rate, the number of assembled reads, assembled contig length, N50, reference coverage, and specificity. iMPP successfully recovered ∼500-2,000 more known protein sequences than the second-best method and reconstructed ∼700-14,000 more novel peptide sequences over 60aa. Taken together, iMPP has demonstrated the feasibility and benefit of integrating *de novo* nucleotide assembly, gene calling, and peptide assembly in metagenomic functional analysis.

## MATERIALS AND METHODS

### The iMPP Algorithm

#### iMPP overview

Figure 2 summarizes the iMPP workflow. iMPP first runs FragGeneScan (42) on the unassembled metagenomic (MG) reads to perform fragmented gene calling. In order to leverage sequence overlap information to improve gene calling, iMPP uses nucleotide assemblers SGA (55) and SPAdes (37) to generate assembly overlap graph and de Bruijn graph contigs, respectively. It then merges them into a hybrid graph (see the “Assembly Graph Merging” section). iMPP performs the second pass of gene calling on the edges and paths of the hybrid graph (see the “iMPP Gene Calling” section). Subsequently, iMPP refines the gene calling results by exploiting sequence overlap information among the peptide reads (see the “Gene Calling Refinement” section). Finally, all predicted short peptides are assembled using PLASS (49). Below we focus on the three modules uniquely contributed by iMPP (Figure 2, bolded operations). More detailed method descriptions, including the chosen parameters and command lines, are available from Supplementary Methods.

**Figure 2:**
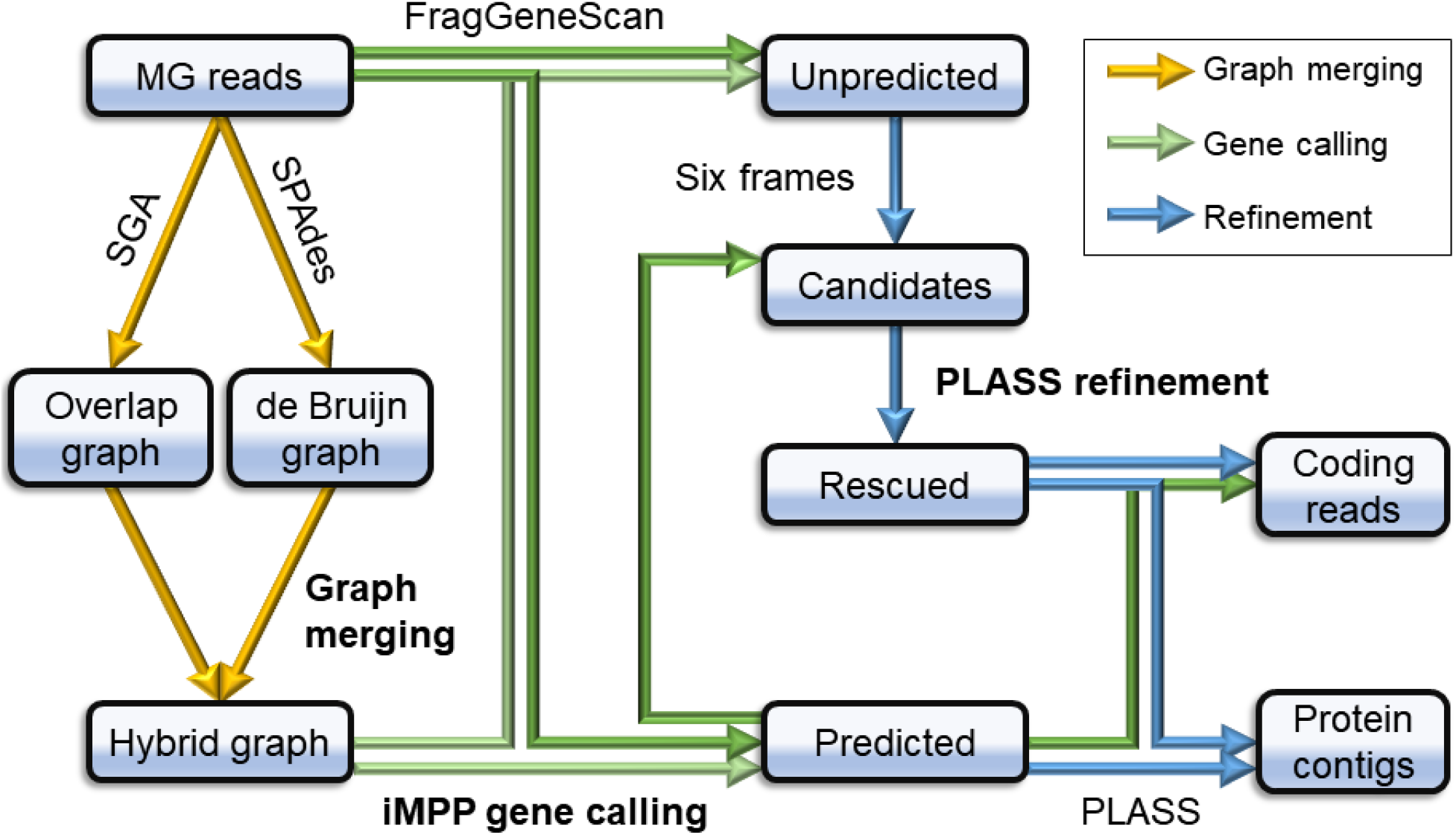
The iMPP workflow overview. Yellow arrows indicate nucleotide assembly information flow; green arrows indicate gene calling information flow; and blue arrows indicate peptide assembly information flow. The bolded operations, i.e., “graph merging”, “iMPP gene calling”, and “PLASS refinement” are unique contributions of iMPP and discussed in detail in the Materials and Methods section. “MG reads” stands for metagenomic reads.

#### Assembly Graph Merging

iMPP employs a hybrid graph generation module that combines a nucleotide assembly overlap graph and a set of contigs generated by de Bruijn graph assemblers (Figure 3A). de Bujin graph assembly breaks down the reads into *k*-mers and models sequence overlap via shared *k*-mers among reads. It can identify sequence overlaps with a greater sensitivity and often produces more complete assemblies. However, it may overlook minor local sequence variations due to its more aggressive graph simplification strategy. Overlap graph, in contrast, preserves raw sequence variation information but is more fragmentary. Therefore, by merging information from both graphs, we expect to preserve the raw sequence information from the overlap graph and improve the graph connectedness. The idea is similar to hybrid assembly, where longer reads (e.g., PacBio SMRT or Oxford Nanopore MinION) are used to connect short reads (56-58) to improve the overall assembly.

**Figure 3:**
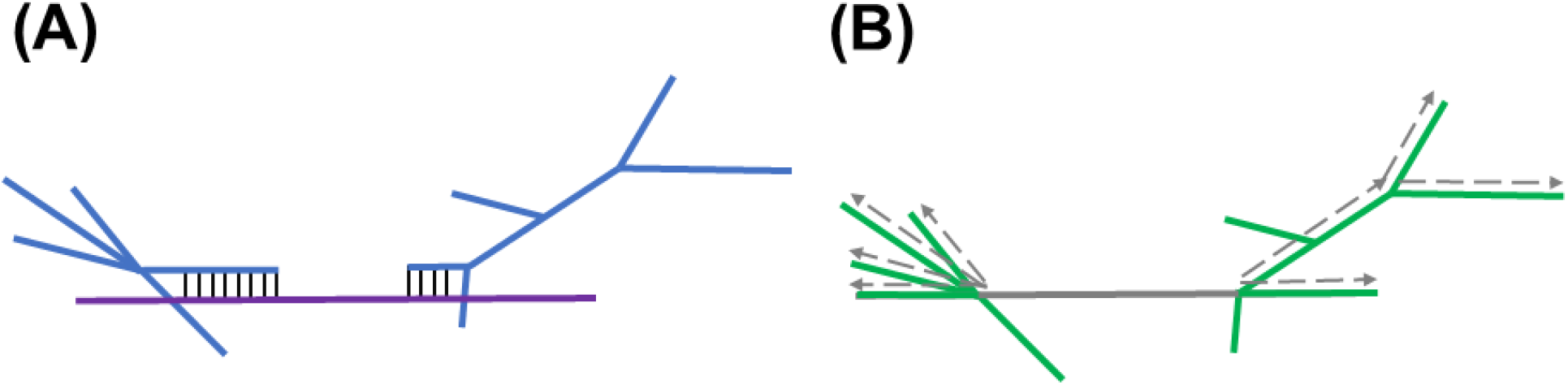
A schematic illustration of the iMPP hybrid graph generation and iMPP gene calling modules. (A) iMPP aligns terminal edges of the assembly overlap graph components (blue) against the de Bruijn graph contigs (purple), and attaches the aligned overlap graph components to the de Bruijn graph contigs. (B) iMPP predicts the protein-coding potential of each edge of the resulted hybrid graph and marks them as either protein-coding (green) or noncoding (gray). iMPP selects the noncoding edges (gray) as anchors and performs depth-first-search from the anchors towards both directions to generate candidate paths.

Specifically, iMPP attempts to connect the isolated overlap graph components using de Bruijn graph contigs as a bridge. iMPP generates an assembly overlap graph using SGA (55) and simplifies the overlap graph by collapsing all unbranched unipaths into single paths (Figure 3A, the blue graph). iMPP then uses SPAdes (under “meta” mode (37)) to generate de Bruijn contigs (Figure 3A, the purple sequence). Denote a vertex with an in- or out-degree of 0 as a *dead end* and an edge containing at least one dead end as a *terminal edge*. iMPP collects all terminal edges from the overlap graph and maps them against all de Bruijn graph contigs. It discards the alignments in which the dead-end sequences are clipped. iMPP then attaches the overlap graph components onto the aligned de Bruijn graph contigs (Figure 3A). iMPP includes all unaligned overlap graph components and de Bruijn graph contigs into the hybrid graph without any modification. This module is similar to the hybrid graph construction module in DRAGoM (59).

#### iMPP Gene Calling

Given the hybrid graph, iMPP performs the second pass of gene calling on the paths of the hybrid graph (recall that the first pass of gene calling is performed directly on unassembled reads). Since paths in the hybrid graph contain sequences longer than individual reads, they may contain more complete and stable ORF signals (26,60). However, as the number of paths grows exponentially w.r.t the traversal depth, iMPP employs an “anchor and extend” heuristic to reduce the running time. Specifically, iMPP first runs FragGeneScan (42) on the edges of the hybrid graph. Since microbial genomes are dense in protein-coding genes, the graph usually contains significantly fewer unpredicted edges (i.e., noncoding) than predicted edges. Consequently, iMPP only selects the unpredicted edges as anchors to avoid traversing a large proportion of the graph (Figure 3B). Intuitively, if many predicted edges surround an unpredicted edge, the unpredicted edge is likely to be protein-coding and should also be predicted. iMPP performs a depth-first search (DFS) towards both directions from each anchor (Figure 3B). The DFS terminates after reaching a certain depth, which further bounds the number of paths that need to be reinvestigated. Finally, iMPP reperforms gene calling on the collected paths using FragGeneScan (42). The predicted edges and paths are both considered as protein-coding; the MG reads that can be mapped to the protein-coding edges and paths are considered as protein-coding reads.

#### Gene Calling Refinement

iMPP further refines the gene calling results by utilizing the overlap information in peptide space. Note the difference between this stage and the previous stage, which relies on overlap information in nucleotide space. Due to codon redundancy, reads that cannot be overlapped in nucleotide space (because of synonymous mutations) may be overlapped in peptide space (47). Hence, ORF signals that are missed during nucleotide assembly could be captured by peptide assembly. Specifically, iMPP collects the remaining unpredicted reads and performs all six-frame translations to convert them into *pseudo peptides*. Note that each nucleotide read can associate with up to six pseudo peptides. Then, the pseudo peptides are assembled with the predicted peptides using PLASS (49). Reads with at least one of their pseudo peptides assembled into long-enough contigs are considered as protein-coding; the pseudo peptides contained in the longest contigs are used to determine the frame.

### Benchmark Datasets

We used four real datasets from different environments (human gut (61), soil (62), marine (63), and cow rumen (64)) to benchmark iMPP. We named them DS1, DS2, DS3, and DS4, respectively. To obtain ground truth, we collected microbial genomes typically found from the corresponding environment and mapped the reads against these reference genomes. The reference genomes and their relative abundances for DS1-4 are available from Supplementary Tables 12-15, respectively. We compiled all the mapped reads into so-called *subsampled datasets*. We also used the entire set of reads to benchmark the software’s performance on real data; we refer to them as the *complete datasets*. Detailed information is summarized in Table 1 and is available in Supplementary Methods.

**Table 1:** A summary of benchmark datasets. “#Reads” indicates the total number of reads for the complete dataset; “#Genomes” indicates the number of reference genomes used for subsampling, “#Sampled Reads” corresponds to the number of subsampled reads.

| Dataset | Accession | Description | Len. | #Reads | #Genomes | #Sampled Reads |
| --- | --- | --- | --- | --- | --- | --- |
| DS1 | SRR341583 | Human Gut | 75 | 23.4M | 3,499 | 9M |
| DS2 | SRR350919 | Soil | 75 | 43.9M | 65,356 | 2.9M |
| DS3 | SRR5720229 | Marine | 150 | 61.1M | 27,216 | 11.3M |
| DS4 | ERR2027889 | Cow Rumen | 150 | 109.8M | 8,980 | 7.8M |

We also benchmarked using three simulated datasets, where the first two comprised of reads generated *in silico* from reference genomes, and the third was a CAMI dataset (65). Please see Supplementary Methods and Results as well as Supplementary Table 2 for more information regarding these simulated datasets.

### Performance Metrics

We benchmarked iMPP in terms of both *de novo* gene calling and peptide assembly. For gene calling, we compared iMPP with three other strategies. The first strategy corresponded to fragmented gene calling directly on unassembled reads using FragGeneScan (42), denoted as “FGS” for short. The second strategy was to assemble the reads using SGA and then performed gene calling using FragGeneScan on the assembled contigs. We denote this strategy as “SGA+FGS”. The third strategy was similar to the second one, but with SPAdes as the assembler, denoted as “SPAdes+FGS”. We measured the performance of iMPP and these three strategies using precision and recall. For the subsampled datasets, we used FragGeneScan to identify all protein-coding regions from the reference genomes. We chose FragGeneScan for ground truth generation because it was also used by the benchmarked strategies (including iMPP), eliminating the impact of using different gene callers. We defined true positives (TP) as the predicted reads with >60% of their total lengths mapped to the coding regions in the reference genomes, false positives (FP) as the predicted reads that are not mapped to the coding regions, and false negatives (FN) as the unpredicted reads that can be mapped to the coding regions. Then, we computed the recall, precision, and F-score as:

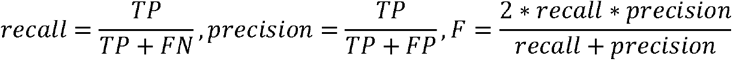

Since no ground truth was available for the complete datasets, we only reported the number of predicted protein-coding reads.

For peptide assembly benchmark, we benchmarked iMPP with two other strategies. The first strategy corresponded to the assembly of FragGeneScan (42) predicted reads using PLASS (49). This strategy was similar to SPA (47,48), which expected the input to be selected by gene callers. We refer to this strategy as “FGS+PLASS”. The second strategy was to use the entire set of unfiltered reads, which was the expected input of PLASS. We refer to this strategy as “PLASS”. We measured the total number of assembled reads, the number of output contigs, the total length of output contigs, N50, assembly rate (the number of assembled reads overall the total number of reads), and chimera rate. To evaluate the correctness of the assembly, we further aligned (using DIAMOND (66)) the contigs against the proteins encoded in the reference genomes (for the subsampled datasets) and the UniProt (31) database (for the complete datasets). A contig was considered true if its aligned proportion was above a certain threshold. We reported *contig-level specificity* as the total length of the true contigs over the total contig length, and *read-level specificity* as the total number of reads constituting the true contigs over the total number of assembled reads. Finally, to measure sensitivity, we reported reference coverage as the percentage of reference genes covered (for more than a length threshold) by the assembled contigs.

## RESULTS

### Gene Calling Benchmark

The gene calling performances of iMPP and the other strategies on the four subsampled datasets are summarized in Figure 4. The results were broadly consistent among all datasets, where iMPP demonstrated the highest performance, followed by FGS, SPAdes+FGS, and SGA+FGS. Specifically, the peak F-scores of iMPP were 92.98%, 92.13%, 92.50%, and 92.73% on the four datasets, respectively (Supplementary Table 1). The second-best strategy, FGS, showed F-scores of 87.22%, 84.72%, 89.54%, and 88.56%, respectively. iMPP improved over FGS with an F-score of 2.96% - 5.76%. Given the already-high performance baseline of >85% F-score, the improvement was significant. Strategies that perform gene calling on assembled reads, i.e., SGA+FGS and SPAdes+FGS, performed worse than FGS, potentially because many reads were not assembled into contigs and were not considered.

**Figure 4:**
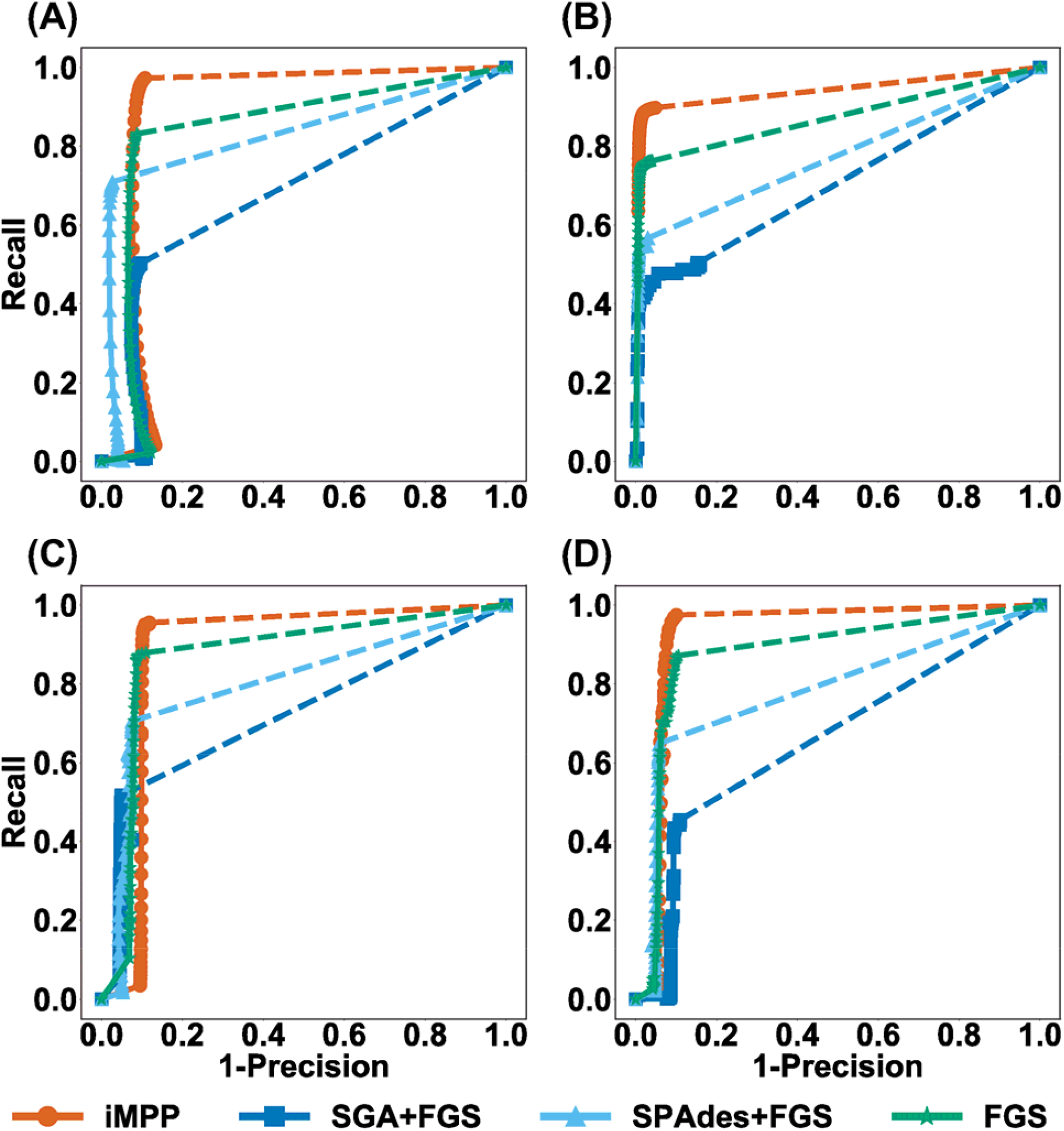
The ROC curves for the *de novo* gene calling performances of the four strategies on the subsampled datasets. (A) DS1, (B) DS2, (C) DS3, and (D) DS4.

For the gene calling performance on the complete datasets, we only report the raw prediction counts because no ground truth is available (Figure 5). iMPP and FGS predicted more protein-coding reads than the assembly-based strategies SGA+FGS and SPAdes+FGS. This is likely due to the low assembly rate on these datasets. iMPP also predicted more reads than FGS, especially on DS1 and DS2 (14.61% and 20.96% more, respectively). The improvement was marginal on DS3 and DS4 (6.71% and 2.89%, respectively). The results are consistent with the observations made from the subsampled datasets (Figure 4), where iMPP showed the highest recall rate among all strategies.

**Figure 5:**
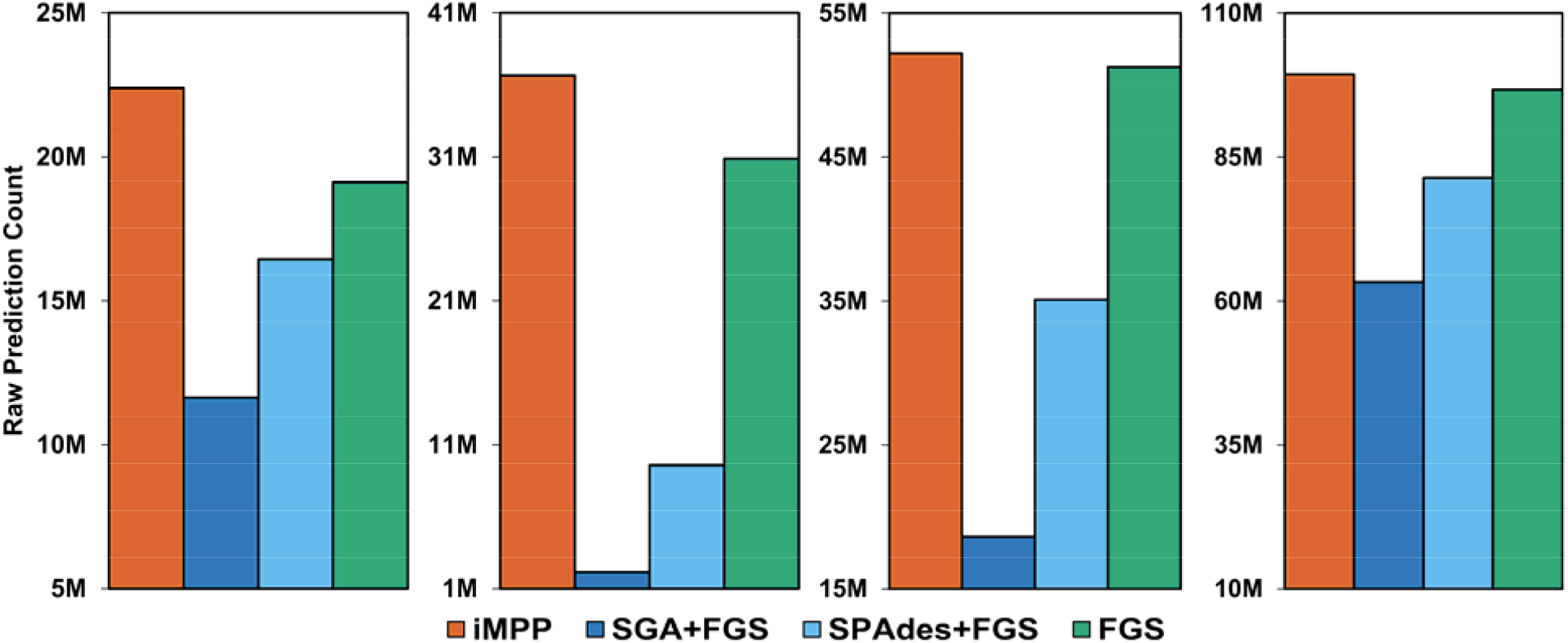
The number of protein-coding reads predicted by iMPP, SGA+FGS, SPAdes+FGS, and FGS on the four complete datasets. Panels from left to right: DS1, DS2, DS3, and DS4.

### Peptide Assembly Benchmark

We summarize the peptide assembly benchmark results on the subsampled datasets in Table 2. We only considered peptide contigs that are ≥60aa long. iMPP assembled the largest number of contigs and total contig length for all datasets. It outperformed FGS+PLASS by 3.3% - 12.5% of the total contig length, but with a less significant improvement over PLASS (0.2% - 4%). Note that the peptide assembly module of iMPP only accepted the predicted protein-coding reads as input, which is less than the entire dataset accepted by the PLASS strategy (DS1: 8.7M vs 9.0M, DS2: 2.4M vs 2.9M, DS3: 11.1M vs 11.3M, and DS4: 7.4M vs 7.8M). However, even with fewer input reads, iMPP assembled more contigs in terms of both the quantity and total length. It suggests that eliminating noncoding reads from the input can potentially benefit peptide assembly. On the other hand, iMPP also outperformed the FGS+PLASS approach that also refined the input. It suggests that true protein-coding reads should not be excluded from the input or it might harm peptide assembly. Because of using the most accurate input sets, iMPP had the highest assembly rate overall. Although slightly underperformed FGS+PLASS on DS1 and DS2 (∼0.3%), iMPP showed a significantly higher assembly rate than FGS+PLASS on DS3 and DS4 (9.5% and 9.3% improvement, respectively). iMPP also consistently showed the highest N50, although it remained similar to the other strategies. Finally, iMPP showed a slightly higher chimera rate, but it remained ignorable and below 0.1% on all datasets.

**Table 2:** Peptide assembly statistics of iMPP, FGS+PLASS, and PLASS on the subsampled datasets. The highest performance in each category is bolded.

| Dataset | Metrics | iMPP | FGS+PLASS | PLASS |
| --- | --- | --- | --- | --- |
| **DS1**<br>**(subsampled)** | # Contigs | **28,645** | 25,621 | 26,833 |
|  | Assembly rate (%) | 26.65 | **26.81** | 24.97 |
|  | Total contig length | **2,180,987** | 1,945,668 | 2,095,045 |
|  | N50 (bp) | **75** | **75** | 74 |
|  | Chimera rate (%) | 0.026 | **0.018** | 0.021 |
| **DS2**<br>**(subsampled)** | # Contigs | **6,911** | 6,030 | 6,680 |
|  | Assembly rate (%) | 7.92 | **8.21** | 6.42 |
|  | Total contig length | **591,348** | 525,637 | 585,397 |
|  | N50 (bp) | **87** | **87** | **87** |
|  | Chimera rate (%) | **0.00** | **0.00** | **0.00** |
| **DS3**<br>**(subsampled)** | # Contigs | **761,837** | 720,195 | 758,779 |
|  | Assembly rate (%) | **60.53** | 51.00 | 58.18 |
|  | Total contig length | **79,683,890** | 77,128,267 | 79,561,338 |
|  | N50 (bp) | **111** | 110 | **111** |
|  | Chimera rate (%) | 0.014 | **0.008** | 0.012 |
| **DS4**<br>**(subsampled)** | # Contigs | **2,591,823** | 2,235,889 | 2,537,281 |
|  | Assembly rate (%) | **91.62** | 82.36 | 87.46 |
|  | Total contig length | **370,451,823** | 350,734,063 | 369,827,910 |
|  | N50 (bp) | **180** | 179 | **180** |
|  | Chimera rate (%) | 0.093 | 0.099 | **0.092** |

We aligned the resulted contigs against the ground-truth reference proteins to investigate the accuracy of the peptide assemblies. The contig- and read-level specificities of different strategies are summarized in Table 3. All three strategies had similar levels of performance, with most of the differences <2%. iMPP showed the highest assembly accuracy in DS1, DS3, and DS4, while FGS+PLASS was the best for DS2. It is expected that iMPP and FGS+PLASS had higher assembly accuracies, as they accepted only the predicted protein-coding reads as input. On the other hand, PLASS used all reads, including noncoding reads, which could have compromised the assembly accuracy.

**Table 3:** The contig- and read-level specificity (%) for the peptide assemblies made by iMPP, FGS+PLASS, and PLASS at different length thresholds on the subsampled datasets. The highest performance in each category is bolded.

| Dataset | Len | iMPP |  | FGS+PLASS |  | PLASS |  |
| --- | --- | --- | --- | --- | --- | --- | --- |
|  |  | contig | read | contig | read | contig | read |
| **DS1**<br>(subsampled) | 60% | **83.62** | **90.07** | 83.46 | 89.93 | 83.55 | 89.72 |
|  | 70% | **83.32** | **89.64** | 83.18 | 89.61 | 83.24 | 89.27 |
|  | 80% | **83.15** | **88.81** | 82.60 | 88.80 | 82.64 | 88.46 |
|  | 90% | **82.86** | **87.21** | 81.24 | 87.14 | 81.36 | 86.81 |
| **DS2**<br>(subsampled) | 60% | 98.61 | 99.45 | **98.74** | **99.70** | 98.58 | 99.68 |
|  | 70% | 98.55 | 99.22 | **98.62** | **99.60** | 98.33 | 99.54 |
|  | 80% | 98.43 | 98.83 | **98.45** | **99.32** | 98.01 | 99.06 |
|  | 90% | **97.83** | 98.47 | 97.49 | **98.50** | 97.23 | 97.86 |
| **DS3**<br>(subsampled) | 60% | **72.82** | **67.49** | 69.90 | 61.36 | 72.03 | 66.24 |
|  | 70% | **72.11** | **65.82** | 68.84 | 58.17 | 71.01 | 63.33 |
|  | 80% | **71.73** | **60.25** | 66.85 | 53.03 | 69.03 | 58.95 |
|  | 90% | **62.28** | 42.44 | 58.87 | 43.88 | 60.82 | **50.99** |
| **DS4**<br>(subsampled) | 60% | **84.00** | **96.81** | 83.60 | 96.89 | 83.00 | 96.16 |
|  | 70% | **83.74** | 96.22 | 83.34 | **96.44** | 82.73 | 95.66 |
|  | 80% | **83.18** | **95.97** | 82.78 | 95.82 | 82.16 | 94.93 |
|  | 90% | **81.59** | **94.80** | 81.03 | 94.64 | 80.58 | 93.70 |

We further calculated the proportion of reference protein sequences recovered by the assemblies generated by difference strategies (Figure 6). iMPP consistently showed the highest reference coverages at all sequence length thresholds on all four benchmark datasets. The average improvement over the second-best PLASS strategy was 1.1%. The results were in line with the observation that iMPP generated more contigs and total contig length than PLASS (Table 2). Taken together, iMPP showed the highest *de novo* peptide assembly sensitivity and accuracy on the subsampled datasets.

**Figure 6:**
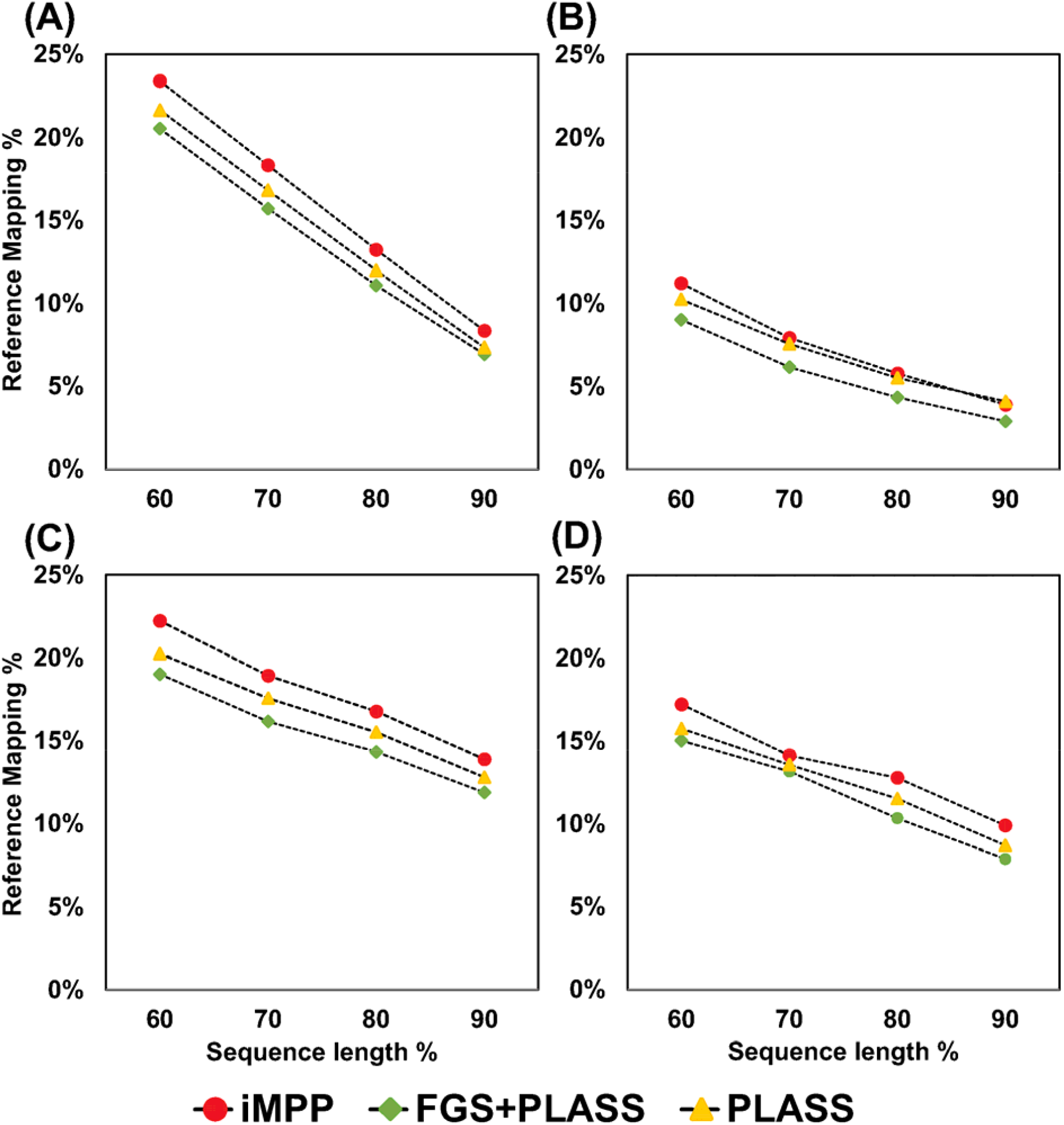
Reference coverages by the peptide contigs assembled by iMPP, FGS, and PLASS on the four subsampled datasets. (A) DS1, (B) DS2, (C) DS3, and (D) DS4.

We also performed similar analyses on the complete datasets. The results summarized in Table 4 were largely consistent with what had been observed for the subsampled datasets (Table 2). Specifically, iMPP assembled significantly more contigs (2.3% - 9.9%) and longer total contig length (1.8% - 10.0%) than FGS+PLASS, and slightly more than PLASS (0.4% - 3.0% more assembled contigs, 0% - 1.2% longer total contig length). The only exception was on DS3, where PLASS assembled slightly more contigs (<0.1%) than iMPP but had shorter total contig length. It is likely that the PLASS assembly was more fragmentary, as indicated by its lower N50 in DS3. Meanwhile, iMPP consistently showed the highest assembly rate and N50 among all datasets, although with marginal improvements (the only exception was that iMPP had a 0.5% lower assembly rate than FGS+PLASS on DS1). All strategies had the same low chimera rate.

**Table 4:** Peptide assembly statistics of iMPP, FGS+PLASS, and PLASS on the complete datasets. The highest performance in each category is bolded.

| **Dataset** | **Metrics** | **iMPP** | **FGS+PLASS** | **PLASS** |
| --- | --- | --- | --- | --- |
| **DS1**<br>**(complete)** | # Contigs | **69,943** | 65,254 | 69,289 |
|  | Assembly rate (%) | 24.65 | **25.18** | 23.21 |
|  | Total contig length | **5,290,265** | 4,933,737 | 5,234,898 |
|  | N50 (bp) | **75** | 74 | 74 |
|  | Chimera rate (%) | **0.00** | **0.00** | **0.00** |
| **DS2**<br>**(complete)** | # Contigs | **515,905** | 469,788 | 510,832 |
|  | Assembly rate (%) | **9.72** | 9.53 | 7.92 |
|  | Total contig length | **55,942,224** | 50,841,297 | 55,692,303 |
|  | N50 (bp) | **111** | 110 | **111** |
|  | Chimera rate (%) | **0.00** | **0.00** | **0.00** |
| **DS3**<br>**(complete)** | # Contigs | 7,325,292 | 7,149,433 | **7,328,392** |
|  | Assembly rate (%) | **41.44** | 39.54 | 35.16 |
|  | Total contig length | **884,928,849** | 869,609,146 | 884,791,359 |
|  | N50 (bp) | **131** | 130 | 130 |
|  | Chimera rate (%) | **0.00** | **0.00** | **0.00** |
| **DS4**<br>**(complete)** | # Contigs | **30,876,275** | 30,179,658 | 30,862,245 |
|  | Assembly rate (%) | **66.16** | 66.02 | 59.64 |
|  | Total contig length | **951,090,439** | 880,119,183 | 950,690,715 |
|  | N50 (bp) | **205** | **205** | **205** |
|  | Chimera rate (%) | **0.00** | **0.00** | **0.00** |

As we did not have the ground truth reference proteins for the complete datasets, we aligned the assembled peptide contigs against the UniProt database (31) to benchmark assembly accuracy. The corresponding contig- and read-level specificities are summarized in Table 5. The results were again consistent with the subsampled datasets, with iMPP leading most of the metrics. All accuracies were lower than those for the subsampled datasets, as the complete dataset may contain more novel proteins that cannot be aligned. Interestingly, the second-most accurate strategy appeared to be PLASS for the complete datasets, unlike FGS+PLASS for the subsampled datasets. The reason could be that the ORF model used by FragGeneScan was trained on known protein families and might miss true protein-coding reads from the novel protein families in the complete datasets. The less complete input further led to fragmentary assemblies, where many short contigs failed to be reliably aligned to references.

**Table 5:** The contig- and read-level specificity (%) for the peptide assemblies made by iMPP, FGS+PLASS, and PLASS at different length thresholds on the complete datasets. The highest performance in each category is bolded.

| Dataset | Len | iMPP |  | FGS+PLASS |  | PLASS |  |
| --- | --- | --- | --- | --- | --- | --- | --- |
|  |  | contig | read | contig | read | contig | read |
| **DS1**<br>(complete) | 60% | **27.43** | **50.89** | 26.81 | 47.71 | 27.12 | 48.11 |
|  | 70% | **27.39** | **50.23** | 26.31 | 46.91 | 27.04 | 48.02 |
|  | 80% | **26.92** | **49.78** | 25.41 | 45.10 | 26.81 | 47.71 |
|  | 90% | **26.54** | **48.25** | 23.30 | 43.26 | 26.31 | 46.21 |
| **DS2**<br>(complete) | 60% | **46.93** | 54.11 | 45.80 | **54.28** | 46.33 | 53.72 |
|  | 70% | **46.25** | **53.67** | 45.11 | 53.46 | 45.69 | 52.97 |
|  | 80% | **44.81** | **52.19** | 43.69 | 51.93 | 44.32 | 51.50 |
|  | 90% | **41.32** | **50.66** | 40.07 | 48.56 | 40.73 | 48.13 |
| **DS3**<br>(complete) | 60% | **44.91** | **54.18** | 43.99 | 52.52 | 44.70 | 53.37 |
|  | 70% | **44.23** | **53.39** | 43.22 | 51.52 | 43.92 | 52.44 |
|  | 80% | **42.74** | **51.72** | 41.46 | 49.53 | 42.14 | 50.60 |
|  | 90% | **38.15** | **48.98** | 35.87 | 44.71 | 36.46 | 46.08 |
| **DS4**<br>(complete) | 60% | **42.35** | **42.28** | 41.73 | 40.76 | 41.76 | 41.24 |
|  | 70% | **41.90** | **41.17** | 41.48 | 40.16 | 41.51 | 40.15 |
|  | 80% | **41.07** | **39.62** | 40.93 | 38.09 | 40.98 | 38.60 |
|  | 90% | **40.63** | **36.78** | 39.46 | 36.18 | 39.53 | 35.75 |

Finally, Figure 7 summarizes the number of peptide contigs that aligned to the UniProt database (31) under different reference length thresholds. iMPP was able to recover the largest number of known protein sequences from UniProt, followed by PLASS (∼500 - 2,000 more peptides). These results reconfirmed iMPP’s high peptide assembly sensitivity on real datasets.

**Figure 7:**
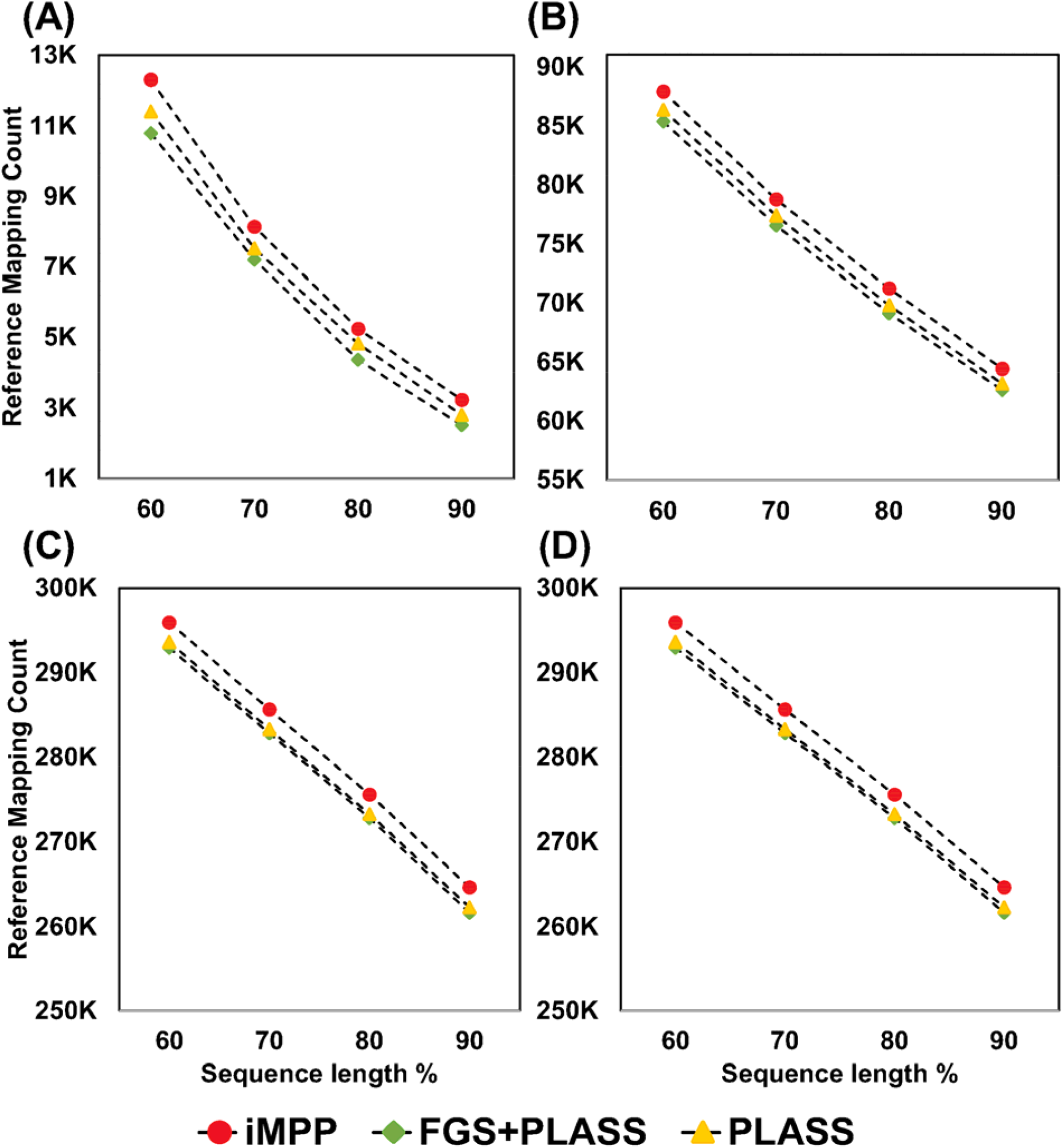
The number of aligned (against UniProt) peptide contigs assembled by iMPP, FGS, and PLASS on the four complete datasets. (A) DS1, (B) DS2, (C) DS3, and (D) DS4.

### Benchmark Results on Simulated Datasets

In addition to the real datasets DS1-DS4, we also benchmarked iMPP on three simulated datasets. We generated two in-house datasets *in silico*, one from 28 marine microbial genomes and one from 8 *Streptococcus* genomes. We also included the CAMI (65) medium-complexity dataset and subsampled it based on the reference genomes provided by the database. For all of the datasets, because of their relatively simple complexity, all methods performed similarly well. More details regarding benchmark results on the simulated datasets can be found from Supplementary Methods and Results as well as Supplementary Figures 1-6 and Supplementary Tables 2-11. The reference genomes used for subsampling and their relative abundances are summarized in Supplementary Tables 16-18.

### Time-Performance Tradeoff

We investigated the proportion of true protein-coding reads discovered by different modules of the iMPP pipeline (Figure 8). Recall that iMPP can make ORF predictions in three stages: from the direct application of FragGeneScan on unassembled reads, from the iMPP gene calling module on the hybrid graph, and finally from the peptide assembly-based refinement. The most economical way to identify coding reads was to perform fragmented gene calling, as FragGeneScan could find >85% of the true positives using ∼10% of the total time. The result was consistent with the high performance observed for FragGeneScan (42). The remaining ∼15% of the protein-coding reads were more challenging to discover, but the majority of them could be discovered using the iMPP gene calling module. It indicates that longer paths from the hybrid assembly graph indeed contain stronger ORF signals and benefit gene calling. However, this module was also the most time-consuming since it performed both overlap graph assembly and de Bruijn graph assembly. It took up ∼55% - 80% of the total runtime of iMPP. Finally, a very small proportion (2% - 3%) of the coding reads could be rescued by peptide assembly-based refinement, which took ∼15% - 30% of the total runtime.

**Figure 8:**
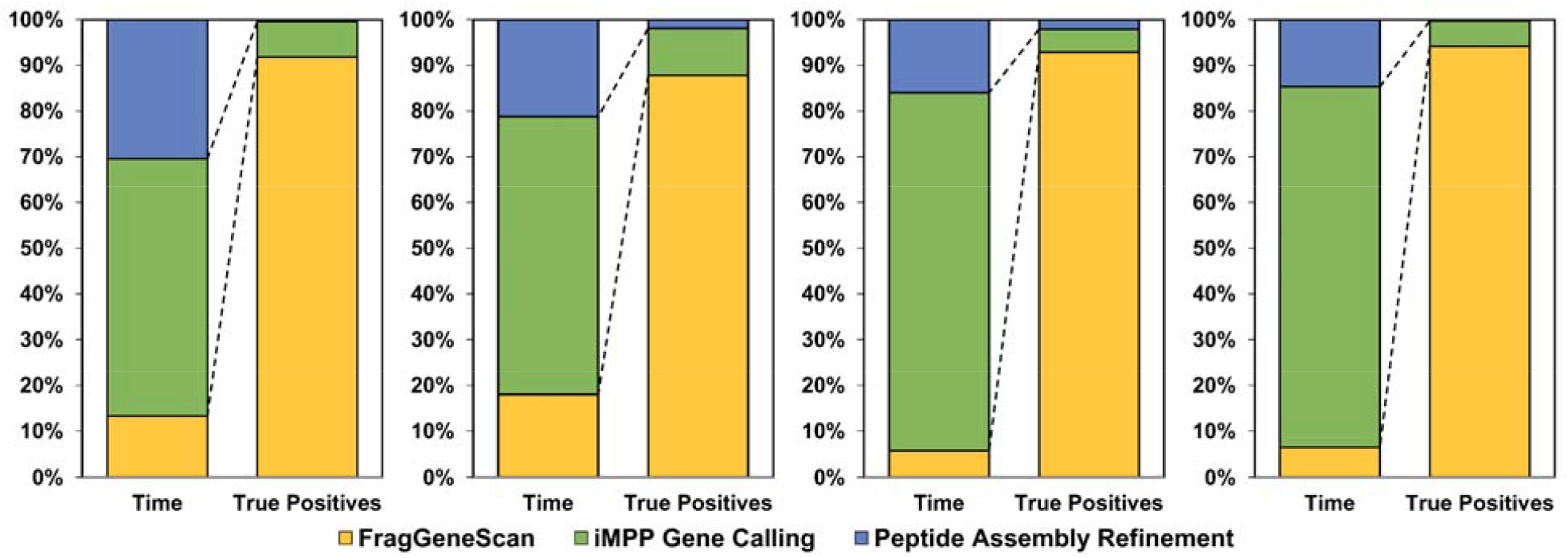
The proportion of the wallclock runtime and detected true protein-coding reads by the direct application of FragGeneScan (yellow), iMPP gene calling (green), and peptide assembly refinement (blue). Panels from left to right: DS1, DS2, DS3, and DS4.

## DISCUSSION

In this work, we present a *de novo* metagenomic functional analysis workflow iMPP. iMPP directly operates on unassembled raw reads and is capable of discovering novel proteins or protein families. To the best of our knowledge, iMPP is currently the only method that integrates nucleotide assembly, gene calling, and peptide assembly based on their informational connection and dependency (Figure 1). The integration appears to be successful based on benchmark results. For gene calling, iMPP significantly improves the state-of-the-art method FragGeneScan with a 10% - 15% higher sensitivity (Figure 4). Notably, while the original sensitivity of FragGeneScan is already high at ∼85%, iMPP has achieved a near-perfect recall rate of >95% at a specificity level of ∼90%. The highly accurate iMPP gene calling results further benefit downstream *de novo* peptide assembly, generating more peptide contigs with higher specificity.

The peptide assembly results point to two seemingly counterintuitive observations regarding *de novo* peptide assembly. First, a more specific input set does not necessarily lead to a more specific assembly. As shown in Figure 4, FragGeneScan had a slightly higher specificity than iMPP (1% - 2%, see details in Supplementary Table 1). However, iMPP assembly showed higher contig- and read-level specificity than FGS+PLASS assembly (Table 3). The reason could be that the more specific input set generated by FragGeneScan was less comprehensive, and missing the true protein-coding reads made the assembly graph more fragmentary. It further resulted in many ultra short peptide contigs, which were subsequently filtered out, reducing the true positive rate and specificity. Second, a more comprehensive input set does not necessarily lead to a more complete assembly. Because iMPP only accepted the predicted coding reads as its input, its input was less complete than PLASS, which accepted all reads. Surprisingly, the iMPP assembly was more comprehensive than the PLASS assembly (Table 2 and Figure 6). The reason could be that contaminants (false pseudo peptides or mispredicted ORF from noncoding reads) may overlap with other peptides by chance, generating more false connections in the assembly graph. The false connections may confound graph traversal and reduce true positive output. As such, a more refined input that contains exactly all the coding reads will likely result in the best assembly. While these observations were made from peptide assembly, we believe that they also apply to nucleotide assembly, as most assembly algorithms are similar. The observation may provide insights to improve *de novo* nucleotide and peptide assembly from a different perspective: refining the input.

We also identified an interesting correlation between environmental characteristics and the difference in assembly performance between subsampled and complete datasets. By comparing Table 2 and Table 4, we found that the assembly rate difference clearly separates the datasets into two subcategories. DS3 (marine) and DS4 (cow rumen) showed a significant increase in assembly rate on the subsampled datasets (20% - 30% improvement) compared to the corresponding complete datasets, whereas DS1 (human gut) and DS2 (soil) showed nearly no improvement (<2%). DS3 and DS4 likely have few known microbial species dominating the corresponding communities. The subsampling process enriched reads from these highly abundant genomes, simplified the subsequent assembly process, and led to a higher assembly rate. Indeed, the most abundant five microbes in DS3 and DS4 comprised 16.1% and 26.1% of the corresponding datasets, respectively (Supplementary Table 14 and 15). On the other hand, the percentage was only ∼2% for DS1 and DS2 (Supplementary Table 12 and 13). In addition to the staggered microbial composition, the completeness of the reference database could also contribute to the smaller assembly rate difference for DS1 and DS2. For DS1 (human gut), we may have identified most of the microbes in the community, such that the subsampled data has similar characteristics as the complete dataset. This is supported by the highest subsampling rate of 38.5% observed in DS1 among all datasets (Table 1). On the other hand, for DS2 (soil), it is possible that our understanding of the environment is so little that the reference-based subsampling process is not significantly better than random subsampling. The conjecture is again in line with the lowest subsampling rate of 6.6% observed in DS2 (Table 1).

With the novel protein families discovered from the above analysis, as well as the known protein families, we expect to develop an algorithm to improve *de novo* nucleotide assembly. This work will complete the last piece of missing information flow from Figure 1 (the gray broken arrow). While it is possible to improve the assembly of individual genes using protein family profiles as a guide (26,54), it is unknown by how much it can improve assembly at the genome level. The improvement observed on individual genes suggests that a guided assembly can help resolve branches in the assembly graph. We shall take advantage of it towards more accurate genome assembly. With this module, we will further develop an iterative version of iMPP following the information flow shown in Figure 1. While the iterative version could be uneconomical given the already high recall rate of the current iMPP version (92% - 97%, Figure 4), it is of theoretical interest to investigate the limit of gene calling directly from fragmented sequences.

To promote practical applications of iMPP, we expect to include an additional module for hypothetical protein annotation. Note that a significant proportion (60% - 70%) of peptide sequences assembled by iMPP from the complete datasets cannot be aligned to the UniProt database (Table 5). Given the high contig-level specificity (∼80%) observed from the subsampled datasets (Table 3), most of the assembled peptide contigs likely correspond to true novel proteins. Note that all of these assembled peptides are ≥60aa, therefore they should contain sufficient information for reliable functional prediction. Specifically, we will develop a hypothetical protein annotation module (67) that includes physicochemical property characterization, domain analysis, protein subcellular localization analysis, and protein-protein interaction analysis. We will also include a *de novo* clustering module to identify novel protein families and sequence motifs (68). Finally, we will also provide the corresponding DNA sequences of these proteins to facilitate their taxonomic analyses and experimental validations.

We also plan to improve the usability and efficiency of iMPP from a software engineering perspective. We will modulate different components of iMPP to meet flexible needs in performance and efficiency. For example, the user will be able to eliminate the peptide assembly refinement step for a speedup without losing a significant number of true positive predictions. We also note that the current iMPP pipeline depends on some third-party software packages, such as SGA (55), SPAdes (37), FragGeneScan (42), and PLASS (49). We will standardize the interface between different modules to allow the substitution of these software packages with other alternatives, e.g., substituting the peptide assembler PLASS (49) with MetaPA (69) or SFA-SPA (48). Finally, we will also try to speed up iMPP by “internalizing” third-party software modules as libraries. For example, iMPP first writes the assembly overlap graph and de Bruijn graph contigs into the hard disk and loads them to generate the hybrid graph. Internalizing the assembly graph generation modules can eliminate the hard-disk traffic and make the entire workflow more efficient. In addition, internalizing peptide assembly will also allow us to access the peptide assembly graph generated by the first PLASS run (for gene calling refinement); the information may help to save a significant amount of time for the second pass (for peptide contig reconstruction).

In conclusion, we present a novel method called iMPP for metagenomic functional analysis. iMPP integrates *de novo* nucleotide assembly, gene calling, and peptide assembly. iMPP is able to improve both gene calling and peptide assembly and has the potential to improve our current understanding of the functions of microbial communities. iMPP was implemented using GNU C++, Python, and Perl. It is freely available from (https://github.com/Sirisha-t/iMPP) under the Creative Commons BY-NC licence.

## Supporting information

Supplementary Methods and Results

