## Supplementary Methods and Results for "Integrated *de novo* Gene Prediction and Peptide Assembly of Metagenomic Sequencing Data"

Sirisha Thippabhotla^1^, Ben Liu^1^, Shibu Yooseph^2^, Youngik Yang^3^, Jun Zhang^4,5^ and Cuncong Zhong^1,^*

^1^Department of Electrical Engineering and Computer Science, The University of Kansas, Lawrence, KS 66045 USA

^2^Department of Computer Science, Genomics and Bioinformatics Cluster, University of Central Florida, Orlando, FL 32816 USA

^3^National Marine Biodiversity Institute of Korea, 101-75, Jangsan-ro, Janghang-eup, Seochun-gun, Chungchungnam-do, 33662, South Korea

^4^Division of Medical Oncology, Department of Internal Medicine, University of Kansas Medical Center, Kansas City, KS 66160 USA

^5^Department of Cancer Biology, University of Kansas Cancer Center; Kansas City, KS 66160 USA

**SUPPLEMENTARY METHODS**

**The iMPP Algorithm**

*Assembly Graph Merging*

Given the input metagenomic sequencing reads, an overlap graph is generated using SGA (v.0.10.15) (1) (sga overlap –m 20), and de Bruijn graph contigs are generated using SPAdes (v.3.9.0) (2) (--only-assembler), with the ‘--meta’ tag enabled for paired-end reads. The SGA overlap graph (.asqg file) is further processed using in-house scripts to condense unbranched paths into single edges. The resulted terminal edges of the SGA overlap graph (i.e., edges containing end nodes that have an in-degree or out-degree of 0) are then aligned with the SPAdes contigs, using BWA-MEM (v.0.7.17) (3). Only those alignments with scores >45 (bwa mem -T 45), alignment length >100, and no clipping at the open end are considered. The overlap graph components are then attached to the aligned SPAdes contigs. SPAdes contigs with no recruited alignment are added as isolated vertices to the hybrid graph. The remaining unaligned overlap graph components are retained without any modification.

*iMPP Gene Calling*

Given the hybrid graph, the fragmentary mode of FragGeneScan (v.1.31) (4) (complete=0 –train=illumina_10) is run against each of the short edges ($\leq$120bp) and the complete mode (complete=1 –train=complete) is run for long edges (>120bp). Edges not predicted by FragGeneScan are considered as anchors. Depth-first search traversal (DFS) on the anchors is performed towards both upstream and downstream, with an extension limit of 150bp on each side. All traversed paths are collected as candidates. Then, the fragmentary mode of FragGeneScan is run on the short candidate paths ($\leq$120bp), and the complete mode is run on the long candidate paths (>120bp). All predicted edges and candidate paths are collected and considered as protein-coding. Then, all reads are mapped against these protein-coding edges and candidate paths using BWA-MEM (3); the alignment score threshold (-T) is set to be 70% of the median length of the input reads.

*Gene Calling Refinement*

The remaining unpredicted reads are collected and translated in all six frames using an in-house script. The translation is terminated when it encounters a stop codon. If it sees a start codon, only the sequence after the start codon is considered. Only the translations that are longer than 20aa are considered. The resulted pseudo peptides are assembled with the predicted reads using PLASS (v.a98349156f9664cac4e3fc7e6df213169a2b82bc) (5) (--min-length 20 --num-iterations 12). Only the peptide contigs that are longer than 60aa are considered. After the assembly, the pseudo peptides are mapped to the resulted peptide contigs using DIAMOND (v.9.24) (6) with default parameters. The reads with at least one of their pseudo peptides mapped (with $\geq$90% sequence identity) are further considered as protein-coding reads. If multiple pseudo peptides are mapped, the frame of the pseudo peptide that can be aligned to the longest peptide contig is considered.

*The other steps*

To perform gene calling directly on the raw reads, FragGeneScan (4) is run in its fragmentary mode (-complete=0 -train=illumina_10). The final peptide assembly stage is performed using PLASS (5) (--min-length 20 --num-iterations 12).

**Benchmark Dataset Generation**

*Real datasets DS1-DS4*

All four real datasets were first quality-trimmed using Trimmomatic (v.0.39) (7) in paired-end mode, with parameters ILLUMINACLIP: TruSeq2-PE.fa:2:30:10 LEADING:3 TRAILING:3 SLIDINGWINDOW:4:15 MINLEN:45*.* After trimming, only the reads that remained paired were retained. To create the subsampled datasets, the reads were then mapped (with default BWA-MEM parameters) to the reference genomes for each dataset. All mapped reads were then collected. The complete list of reference genomes used to subsample DS1, DS2, DS3, and DS4 are available from Supplementary Tables 12, 13, 14, and 15, respectively.

*Simulated datasets SDS1-SDS3*

We also benchmarked iMPP on three simulated datasets (*Streptococcus*, Marine, and CAMI). A summary of the statistics of these datasets can be found in Supplementary Table 2. The *Streptococcus* (SDS1) and Marine (SDS2) datasets were generated using WGSIM (v.0.3.1) (8) ( ‘-1 100 -e 0.01 -r 0’ options) with varying coverages of 2X, 5X, and 10X. The third dataset (SDS3) was downloaded from CAMI (9). We subsampled (using the same subsampling approach as described in the previous section) the dataset to focus on the low-coverage proportion (<10X). The complete list of reference genomes used to subsample SDS1, SDS2, and SDS3 are available from Supplementary Tables 16, 17, and 18, respectively.

**Ground Truth Generation**

The protein-coding regions of the reference genomes were predicted using the complete mode of FragGeneScan (4) (-complete=1, -train=complete). The reads were mapped to the protein-coding regions using BWA-MEM (3); the alignment score threshold (-T) was set to be 70% of the median length of the input reads. The mapped reads were considered to be protein-coding; the other unmapped reads were considered to be non-coding.

**Benchmark Experiments**

*Gene calling benchmark experiment*

For the FGS approach, FragGeneScan was run in its fragmentary mode (-complete=0 -train=illumina_10). For the SGA+FGS approach, SGA was run with parameters “sga assemble -m 20 --transitive-reduction -g 0”, followed by the complete mode of FragGeneScan (-complete=1 -train=complete) on the resulted contigs. For the SPAdes+FGS approach, SPAdes was run with the default parameters, followed by the complete mode of FragGeneScan on the resulted contigs. For all approaches, a range of FragGeneScan score cutoffs (1.10 - 1.50) was used to reveal performances under different stringency levels (as ROC curves).

*Peptide assembly benchmark experiment*

For the FGS+PLASS approach, FragGeneScan was run in its fragmentary mode (-complete=0 -train=illumina_10). The resulted predicted reads were further assembled using PLASS (--min-length 20 --num-iterations 12). For the PLASS approach, all input reads were assembled using PLASS (--min-length 20 --num-iterations 12).

**SUPPLEMENTARY RESULTS**

**The Real Datasets (DS1 - DS4)**

The peak gene-calling performance of iMPP and the other strategies FGS, SPAdes+FGS, SGA+FGS is summarized in Supplementary Table 1.

**The Simulated Datasets (SDS1 - SDS3)**

*Gene calling*

The gene calling performance of iMPP and the other three strategies (FGS, SPAdes+FGS, SGA+FGS) on the simulated datasets SDS1, SDS2, and SDS3 are shown in Supplementary Figures 1, 2, and 3, respectively. The corresponding peak performances are shown in Supplementary Tables 3, 4, and 5, respectively. Because the simulated datasets are relatively simpler than the real dataset, all methods performed well, and no significant difference was observed. iMPP appeared to perform better with low coverage data (2X and 5X coverage on both SDS1 and SDS2), showing a 1-2% improvement of F-score over the second-best strategy FGS (Supplementary Table 3 and 4). With a 10x coverage, the SPAdes+FGS strategy showed the best performance. SPAdes was able to generate near-perfect assemblies at this coverage for the low-complexity simulated datasets, hence lifting the overall performance. iMPP was marginally worse (<0.4%) than SPAdes+FGS on these datasets. iMPP also showed the highest F-score on the CAMI dataset SDS3 (0.3% higher than the second-best method FGS, Supplementary Table 5).

*Peptide assembly*

The peptide assembly statistics on the simulated datasets SDS1, SDS2, and SDS3 are presented in Supplementary Tables 6, 7, and 8, respectively. Similar to the observations made on the real datasets DS1-DS4, iMPP had the highest number of assembled contigs and total contig length for all three datasets. iMPP outperformed FGS+PLASS by 8.92-17.13% of the total contig length and PLASS by a lower percentage improvement of 1.22-3.06%. We also observed a consistent improvement in N50 of iMPP, although marginal. The contig and real-level specificity results on the simulated datasets SDS1, SDS2, and SDS3 are summarized in Supplementary Tables 9, 10, and 11, respectively. All three strategies have similar performance levels, with FGS+PLASS consistently showing the highest specificity at both the contig- and read-level. These reference coverage proportions for SDS1, SDS2, and SDS3 are shown in Supplementary Figures 4, 5, and 6, respectively. Again, all strategies showed similar performance, with iMPP consistently being the best (~0.2-0.45% improvement).

**SUPPLEMENTARY DATA AVAILABILITY**

The complete datasets DS1-4 can be downloaded from NCBI using their SRA accession numbers provided in the manuscript. All other datasets, including subsampled datasets (DS1-4) and those generated in-house (i.e., Supplementary Datasets SDS1-3), can be downloaded from <https://cbb.ittc.ku.edu/iMPP.html>
