## Supplementary Figure for "Integrated *de novo* Gene Prediction and Peptide Assembly of Metagenomic Sequencing Data"

**Supplementary Figures for “Integrated *de novo* Gene Prediction and Peptide Assembly of Metagenomic Sequencing Data”**

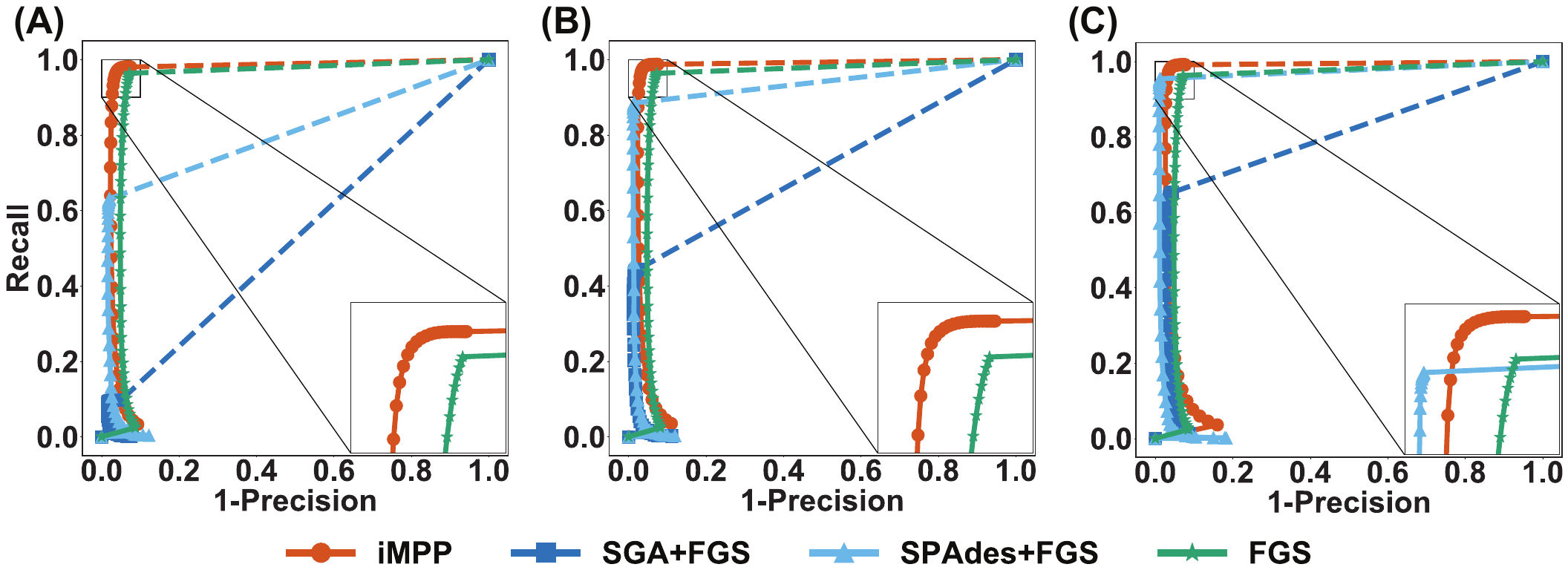

Supplementary Figure 1: The gene prediction performance of different methods on the SDS1 dataset. Figures A, B, C represent performance on SDS1 with 2X, 5X, and 10X coverage, respectively.

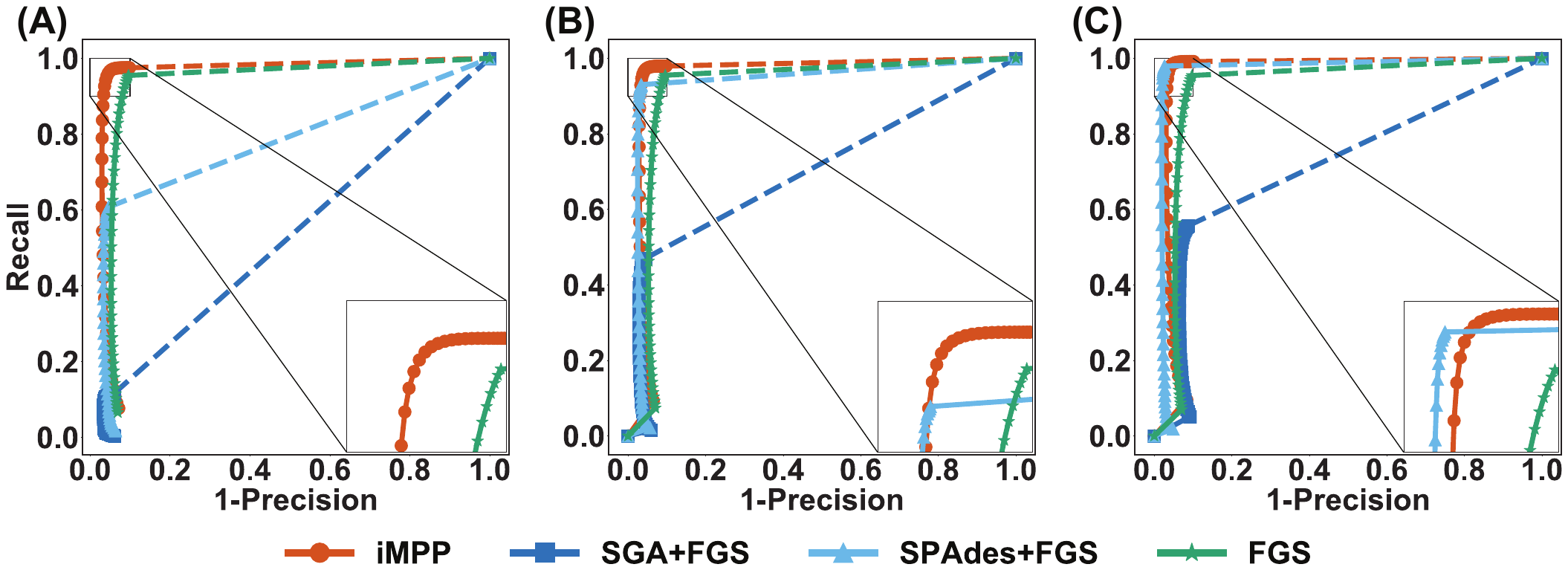

Supplementary Figure 2: The gene prediction performance of different methods on the SDS2 dataset. Figures A, B, C represent performance on SDS2 with 2X, 5X, and 10X coverage, respectively.

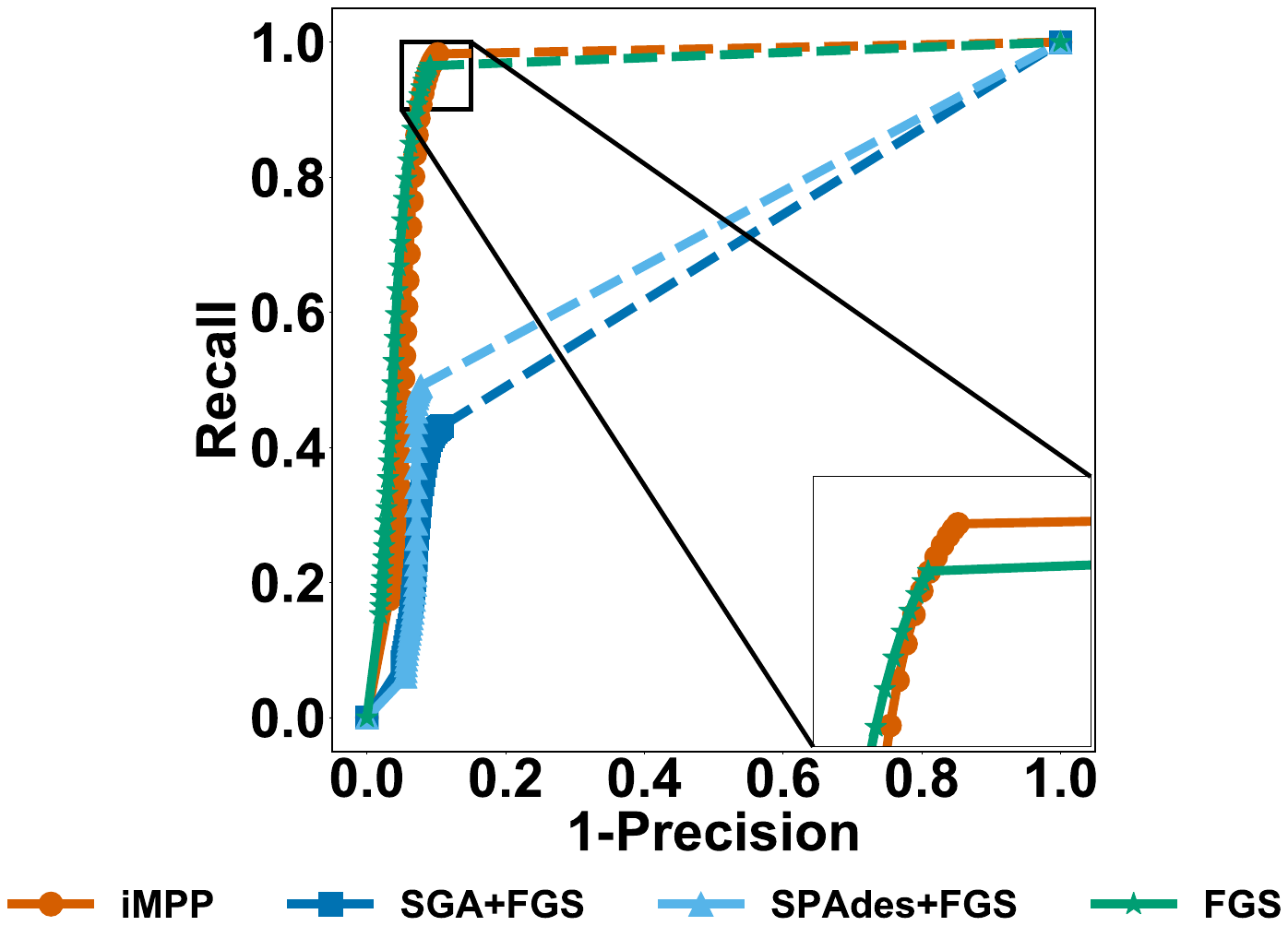

Supplementary Figure 3: The gene prediction performance of different methods on the SDS3 dataset.

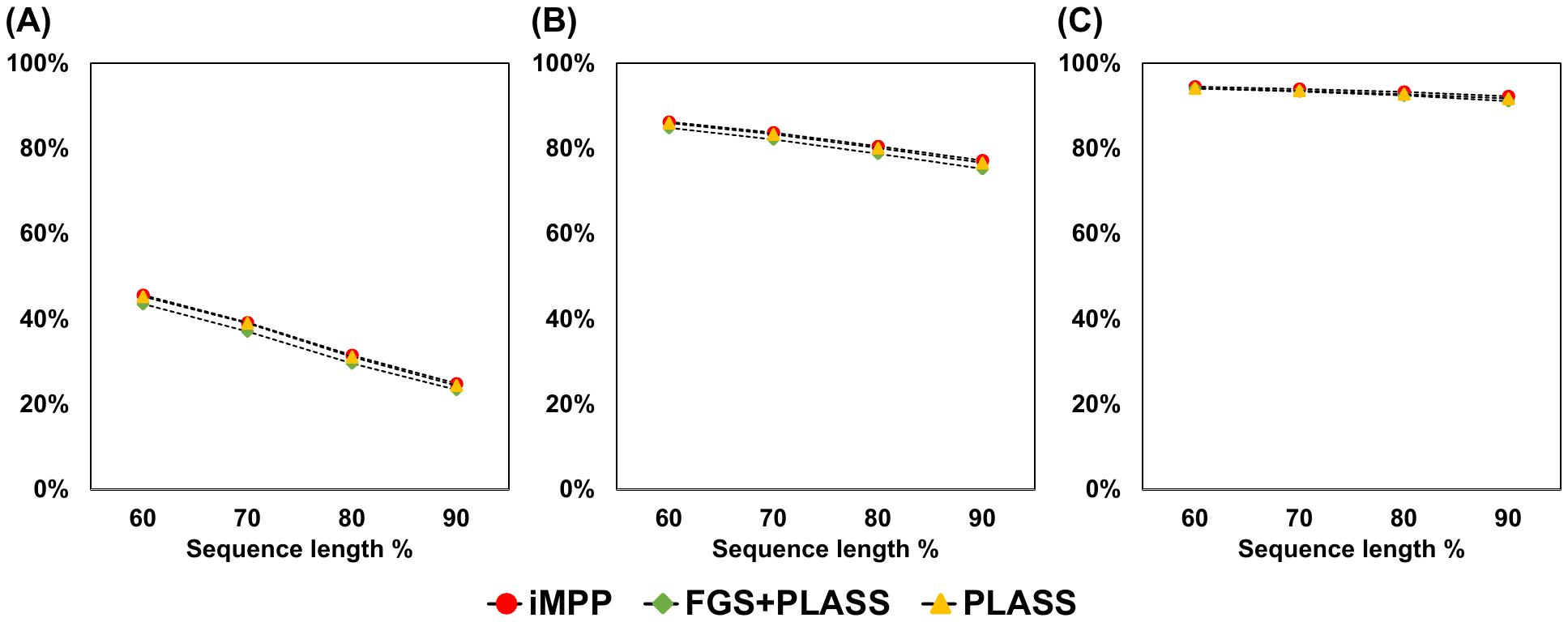

Supplementary Figure 4: Reference coverages by the peptide contigs assembled by iMPP, FGS, and PLASS on simulated dataset SDS1. (A) 2x coverage, (B) 5x coverage, (C) 10x coverage.

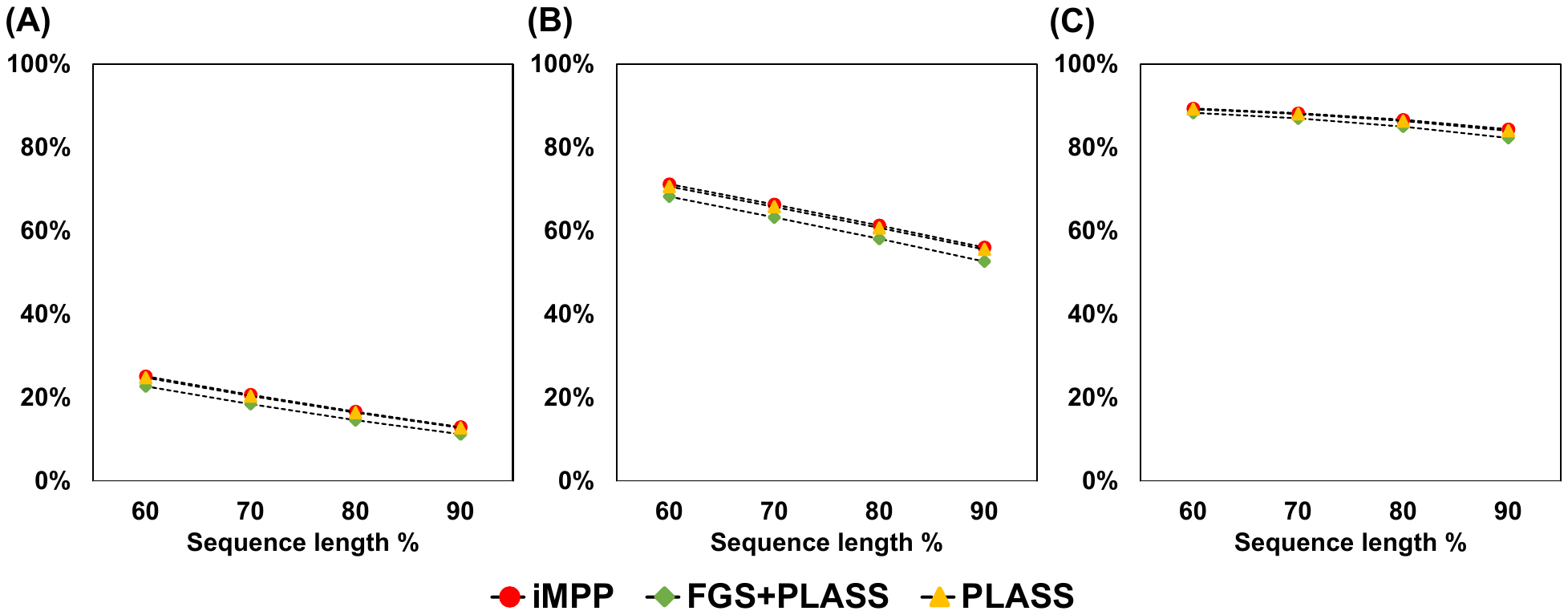

Supplementary Figure 5 : Reference coverages by the peptide contigs assembled by iMPP, FGS, and PLASS on simulated dataset SDS2. (A) 2x coverage, (B) 5x coverage, (C) 10x coverage.

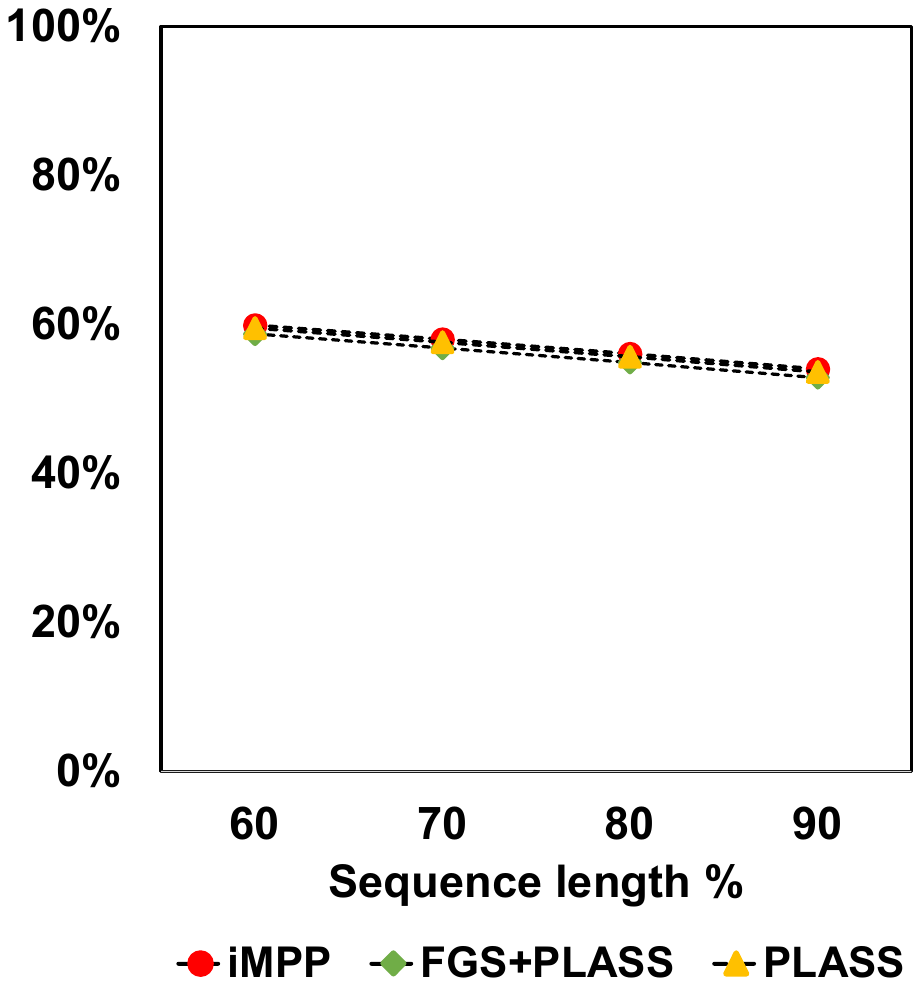

Supplementary Figure 6: Reference coverages by the peptide contigs assembled by iMPP, FGS, and PLASS on simulated dataset SDS3.
