## Supplementary Table for "Integrated *de novo* Gene Prediction and Peptide Assembly of Metagenomic Sequencing Data"

**Supplementary Tables for “Integrated *de novo* Gene Prediction and Peptide Assembly of Metagenomic Sequencing Data”**

Supplementary Table 1: A summary of the optimal (in terms of F-score, among all tested stringency cutoffs) gene prediction performance of different methods on subsampled datasets (DS1-DS4). The best performance within each category is bolded.

| Dataset | Metrics | iMPP | SGA+FGS | SPAdes+FGS | FGS |
| --- | --- | --- | --- | --- | --- |
| DS1  (subsampled) | Precision | 88.94 | 90.00 | **96.99** | 90.71 |
|  | Recall | **97.42** | 50.40 | 71.35 | 84.00 |
|  | F-Score | **92.98** | 64.62 | 82.22 | 87.22 |
| DS2  (subsampled) | Precision | 94.20 | 84.28 | **95.66** | 95.16 |
|  | Recall | **90.16** | 49.12 | 57.15 | 76.35 |
|  | F-Score | **92.13** | 62.06 | 71.55 | 84.72 |
| DS3  (subsampled) | Precision | 89.62 | 94.26 | **95.87** | 91.23 |
|  | Recall | **95.58** | 51.88 | 70.52 | 87.91 |
|  | F-Score | **92.50** | 70.52 | 81.26 | 89.54 |
| DS4  (subsampled) | Precision | 88.66 | 90.46 | **94.03** | 89.40 |
|  | Recall | **97.19** | 45.14 | 64.41 | 87.73 |
|  | F-Score | **92.73** | 60.23 | 76.45 | 88.56 |

Supplementary Table 2: A summary of benchmark simulated datasets. SDS1 and SDS2 have three versions (2X, 5X, 10X).

| Dataset | Description | #Genomes | Error | Read Len. | #Reads |
| --- | --- | --- | --- | --- | --- |
| SDS1 | Streptococcus | 8 | 0.1% | 100 | 0.2M, 0.6M, 1.2M |
| SDS2 | Marine | 28 | 0.1% | 100 | 1.4M, 3.5M, 7M |
| SDS3 | CAMI | 4,679 | - | 100 | 31.3M |

Supplementary Table 3: A summary of the optimal (in terms of F-score, among all tested stringency cutoffs) gene prediction performance of different methods on simulated dataset SDS1, with varying coverages of 2x, 5x and 10x. The best performance within each category is bolded.

| Dataset | Metrics | iMPP | SGA+FGS | SPAdes+FGS | FGS |
| --- | --- | --- | --- | --- | --- |
| SDS1  (2x) | Precision | 91.60 | 89.71 | **96.54** | 92.57 |
|  | Recall | **98.84** | 68.29 | 67.29 | 95.43 |
|  | F-Score | **95.09** | 77.55 | 79.30 | 93.98 |
| SDS1  (5x) | Precision | 91.17 | 89.96 | **97.01** | 92.55 |
|  | Recall | **99.73** | 71.32 | 87.88 | 95.40 |
|  | F-Score | **95.26** | 79.56 | 92.22 | 93.95 |
| SDS1  (10x) | Precision | 90.45 | 91.18 | **97.27** | 92.58 |
|  | Recall | **99.91** | 77.94 | 92.81 | 95.39 |
|  | F-Score | 94.95 | 84.40 | **94.99** | 93.96 |

Supplementary Table 4: A summary of the optimal (in terms of F-score, among all tested stringency cutoffs) gene prediction performance of different methods on simulated datasets SDS2, with varying coverages of 2x, 5x and 10x. The best performance within each category is bolded.

| Dataset | Metrics | iMPP | SGA+FGS | SPAdes+FGS | FGS |
| --- | --- | --- | --- | --- | --- |
| SDS2  (2x) | Precision | 88.78 | 87.23 | **94.17** | 90.58 |
|  | Recall | **98.73** | 69.81 | 65.19 | 93.77 |
|  | F-Score | **93.49** | 77.55 | 77.04 | 92.15 |
| SDS2  (5x) | Precision | 88.78 | 87.23 | **94.17** | 90.58 |
|  | Recall | **99.70** | 73.16 | 91.96 | 93.80 |
|  | F-Score | **93.73** | 80.22 | 92.98 | 92.17 |
| SDS2  (10x) | Precision | 87.92 | 89.98 | **94.15** | 90.48 |
|  | Recall | **99.92** | 78.83 | 93.66 | 94.01 |
|  | F-Score | 93.54 | 84.04 | **93.90** | 92.21 |

Supplementary Table 5: A summary of the optimal (in terms of F-score, among all tested stringency cutoffs) gene prediction performance of different methods on simulated dataset SDS3. The best performance within each category is bolded.

| Dataset | Metrics | iMPP | SGA+FGS | SPAdes+FGS | FGS |
| --- | --- | --- | --- | --- | --- |
| SDS3 | Precision | 89.78 | 89.13 | **92.19** | 90.85 |
|  | Recall | **98.25** | 43.14 | 49.09 | 96.50 |
|  | F-Score | **93.82** | 58.14 | 64.07 | 93.59 |

Supplementary Table 6: Assembly statistics of the iMPP, FGS+PLASS, and PLASS approaches on simulated dataset SDS1 (2x, 5x and 10x coverages), at minimum 60aa length cutoff.

| Dataset | Metrics | iMPP | FGS+PLASS | PLASS |
| --- | --- | --- | --- | --- |
| SDS1  (2x) | # Contigs | **13,897** | 12,572 | 13,678 |
|  | Assembly rate (%) | **25.68** | 24.83 | 24.06 |
|  | Total contig length | **1,328,915** | 1,200,461 | 1,312,935 |
|  | N50 (bp) | **95** | 94 | **95** |
|  | Chimera rate (%) | **0.007** | 0.00 | 0.00 |
| SDS1  (5x) | # Contigs | **72,398** | 66,523 | 71,075 |
|  | Assembly rate (%) | 54.53 | **55.31** | 52.31 |
|  | # Assembled reads | **316,857** | 303,666 | 313,876 |
|  | Total contig length | **8,543,134** | 7,772,851 | 8,306,700 |
|  | N50 (bp) | **123** | 121 | 121 |
|  | Chimera rate (%) | **0.006** | 0.004 | **0.006** |
| SDS1  (10x) | # Contigs | **177,806** | 165,164 | 174,592 |
|  | Assembly rate (%) | 69.91 | **73.27** | 68.74 |
|  | # Assembled reads | **829,222** | 804,336 | 824,925 |
|  | Total contig length | **24,825,269** | 22,791,574 | 24,170,161 |
|  | N50 (bp) | **153** | 151 | 151 |
|  | Chimera rate (%) | 0.023 | 0.021 | **0.035** |

Supplementary Table 7: Assembly statistics of the iMPP, FGS+PLASS, and PLASS approaches on simulated dataset SDS2 (2x, 5x and 10x coverages), at minimum 60aa length cutoff.

| Dataset | Metrics | iMPP | FGS+PLASS | PLASS |
| --- | --- | --- | --- | --- |
| SDS2  (2x) | # Contigs | **35,466** | 30,320 | 34,820 |
|  | Assembly rate (%) | **12.77** | 11.99 | 12.20 |
|  | Total contig length | **3,102,141** | 2,648,529 | 3,055,781 |
|  | N50 (bp) | **84** | **84** | **84** |
|  | Chimera rate (%) | **0.00** | 0.006 | **0.00** |
| SDS2  (5x) | # Contigs | **277,076** | 244,184 | 270,137 |
|  | Assembly rate (%) | **39.37** | 39.22 | 38.51 |
|  | Total contig length | **28,105,620** | 24,354,276 | 27,271,835 |
|  | N50 (bp) | **101** | 99 | **101** |
|  | Chimera rate (%) | **0.003** | 0.005 | 0.005 |
| SDS2  (10x) | # Contigs | **948,115** | 851,645 | 937,977 |
|  | Assembly rate (%) | 64.56 | **66.33** | 64.22 |
|  | Total contig length | **122,752,947** | 107,123,125 | 120,746,454 |
|  | N50 (bp) | **137** | 133 | 136 |
|  | Chimera rate (%) | 0.003 | **0.002** | 0.003 |

Supplementary Table 8: Assembly statistics of the iMPP, FGS+PLASS, and PLASS approaches on simulated dataset SDS3, at minimum 60aa length cutoff.

| Dataset | Metrics | iMPP | FGS+PLASS | PLASS |
| --- | --- | --- | --- | --- |
| SDS3 | # Contigs | **1,673,780** | 1,554,030 | 1,671,639 |
|  | Assembly rate (%) | 70.77 | **72.79** | 69.89 |
|  | Total contig length | **233,415,056** | 215,089,223 | 233,252,311 |
|  | N50 (bp) | **152** | 151 | **152** |
|  | Chimera rate (%) | 0.105 | **0.095** | 0.099 |

Supplementary Table 9: The contig- and read-level specificity (%) for the peptide assemblies made by iMPP, FGS+PLASS, and PLASS at different length thresholds on the simulated dataset SDS1 (2x, 5x and 10x coverages). The highest performance in each category is bolded.

| Dataset |  | iMPP | | FGS+PLASS | | PLASS | |
| --- | --- | --- | --- | --- | --- | --- | --- |
|  | **Seqlen** | **contig** | **read** | **contig** | **read** | **contig** | **read** |
| SDS1  (2x) | 60% | 96.41 | 98.18 | **96.65** | **98.67** | 96.38 | 98.12 |
|  | 70% | 96.24 | 97.88 | **96.49** | **98.43** | 96.22 | 97.84 |
|  | 80% | 95.82 | 97.31 | **96.17** | **97.89** | 95.80 | 97.15 |
|  | 90% | 94.58 | 95.97 | **94.66** | **96.22** | 94.47 | 95.87 |
| SDS1  (5x) | 60% | **93.66** | 97.58 | 93.60 | **98.08** | **93.72** | 97.63 |
|  | 70% | 93.38 | 97.15 | 93.40 | **97.71** | **93.48** | 97.22 |
|  | 80% | 92.88 | 96.50 | **93.06** | **97.02** | 92.97 | 96.57 |
|  | 90% | 91.25 | 94.79 | **92.24** | **95.33** | 91.36 | 94.95 |
| SDS1  (10x) | 60% | 90.70 | 97.09 | **93.60** | **97.76** | 85.67 | 97.07 |
|  | 70% | **90.34** | 96.73 | 87.09 | **97.43** | 86.44 | 96.69 |
|  | 80% | **89.57** | 96.03 | 87.51 | **96.81** | 86.75 | 96.00 |
|  | 90% | **87.23** | 94.25 | 86.24 | **95.15** | 85.60 | 94.30 |

Supplementary Table 10: The contig- and read-level specificity (%) for the peptide assemblies made by iMPP, FGS+PLASS, and PLASS at different length thresholds on the simulated dataset SDS2 (2x, 5x and 10x coverages). The highest performance in each category is bolded.

| Dataset |  | iMPP | | FGS+PLASS | | PLASS | |
| --- | --- | --- | --- | --- | --- | --- | --- |
|  | **Seqlen** | **contig** | **read** | **contig** | **read** | **contig** | **read** |
| SDS2  (2x) | 60% | 95.58 | 97.38 | **96.14** | **97.74** | 95.70 | 97.36 |
|  | 70% | 95.25 | 96.90 | **95.85** | **97.34** | 95.38 | 96.94 |
|  | 80% | 94.75 | 96.27 | **95.40** | **96.79** | 94.83 | 96.18 |
|  | 90% | 93.08 | 94.67 | **93.69** | **95.40** | 93.34 | 94.73 |
| SDS2  (5x) | 60% | 93.92 | 96.48 | **94.76** | **97.08** | 94.04 | 96.51 |
|  | 70% | 93.59 | 95.99 | **94.48** | **96.71** | 93.69 | 96.02 |
|  | 80% | 92.93 | 95.10 | **93.86** | **95.97** | 92.99 | 95.15 |
|  | 90% | 91.01 | 93.20 | **92.03** | **94.13** | 91.12 | 93.23 |
| SDS2  (10x) | 60% | 91.98 | 96.13 | **93.12** | **96.80** | 91.97 | 96.14 |
|  | 70% | 91.50 | 95.61 | **92.74** | **96.36** | 91.48 | 95.59 |
|  | 80% | 90.46 | 94.55 | **91.85** | **95.51** | 90.44 | 94.55 |
|  | 90% | 87.63 | 92.19 | **89.29** | **93.29** | 87.71 | 92.23 |

Supplementary Table 11: The contig- and read-level specificity (%) for the peptide assemblies made by iMPP, FGS+PLASS, and PLASS at different length thresholds on the simulated dataset SDS3. The highest performance in each category is bolded.

| Dataset |  | iMPP | | FGS+PLASS | | PLASS | |
| --- | --- | --- | --- | --- | --- | --- | --- |
|  | **Seqlen** | **contig** | **read** | **contig** | **read** | **contig** | **read** |
| SDS3 | 60% | 88.56 | 96.03 | **89.90** | **96.90** | 88.54 | 95.99 |
|  | 70% | 87.77 | 95.43 | **89.25** | **96.43** | 87.73 | 95.39 |
|  | 80% | 86.08 | 94.38 | **87.78** | **95.53** | 86.06 | 94.36 |
|  | 90% | 81.83 | 92.10 | **83.63** | **93.43** | 81.79 | 92.11 |
